# PqiABC forms a membrane-bridging conduit and mediates bidirectional phospholipid transport across the Gram-negative bacterial envelope

**DOI:** 10.64898/2026.09.15.751761

**Authors:** Hannah E. Johnston, Luke A. Clifton, Charlotte B. Wilson, Pooja Sridhar, Abdulaziz Alzahrani, Adam Colyer, Richard Logan, Stephen C.L. Hall, David J. Hardy, Timothy J. Knowles

## Abstract

The mechanism by which glycerophospholipids are transported between the inner and outer membranes in Gram-negative bacteria remains poorly understood. In *Escherichia coli*, the paraquat-inducible (Pqi) pathway, comprising the inner membrane protein PqiA, the periplasm-spanning MCE-family protein PqiB, and the outer membrane lipoprotein PqiC, has been implicated in this process. These components are proposed to assemble into a quaternary complex that forms a continuous channel bridging the inner and outer membranes.

Here, using neutron reflectometry and quartz crystal microbalance with dissipation monitoring, we perform a dynamic structural analysis of PqiABC within a planar double bilayer membrane-mimetic system. This approach reveals that PqiABC assembles into a stable, envelope-spanning complex anchored to both membranes, consistent with its proposed conduit architecture.

Furthermore, using neutron reflectometry in combination with complementary fluorescence-based assays, we demonstrate that PqiABC mediates passive glycerophospholipid transport, supporting bidirectional lipid exchange between membranes. Together, these findings establish PqiABC as a membrane-bridging lipid transport system and provide direct evidence for a mechanism of passive glycerophospholipid equilibration across the bacterial envelope.

## Introduction

Gram-negative pathogens pose a major and growing global health threat in the era of antibiotic resistance. Their double-membraned cell envelope constitutes a formidable defensive barrier, particularly the outer membrane (OM), which forms a highly impermeable barrier to many molecules, including antibiotics. Disruption of the OM markedly increases antibiotic susceptibility, making its biogenesis a key focus for antimicrobial development. A detailed insight into OM biogenesis may therefore reveal new opportunities for therapeutic intervention.

The OM is an asymmetric bilayer composed primarily of lipopolysaccharide (LPS) in the outer leaflet and glycerophospholipids (GPLs) in the inner leaflet, with a plethora of proteins tightly controlling membrane integrity and trafficking. The pathways responsible for transporting LPS and outer membrane proteins (OMPs) have been extensively characterised (Rollauer et al. 2015, Wilson and Ruiz 2021). In contrast, the transport of GPLs, which are synthesised in the cytosol and at the inner membrane (IM), remains poorly understood.

In recent years, several putative GPL transport systems have been identified in *Escherichia coli* (Yeow and Chng 2022). These include the maintenance of outer membrane lipid asymmetry (Mla) pathway, the lipophilic envelope-spanning tunnel (Let) complex, and the paraquat-inducible (Pqi) complex, all of which contain proteins of the mammalian cell entry (MCE) domain-containing family (Ekiert et al. 2017, Isom et al. 2017). Additional candidates include members of the AsmA-like protein family, which are proposed to form hydrophobic bridges across the cell envelope (Kumar and Ruiz 2023), and the Tol–Pal system (Tan and Chng 2025). Among these, the Mla pathway is the best characterised. It is powered by ATP hydrolysis at the IM by the MlaFEDB complex and mediates retrograde transport of mislocalised GPLs from the OM to the IM via the soluble periplasmic carrier MlaC (Malinverni and Silhavy 2009, Tang et al. 2021, Wotherspoon et al. 2024). In contrast, the Pqi and Let systems contain large, envelope-spanning MCE-domain containing proteins, PqiB and LetB, which assemble into elongated structures with central hydrophobic tunnels. These architectures are proposed to facilitate the direct translocation of hydrophobic cargo, most likely GPLs, across the periplasm (Ekiert et al. 2017). Supporting this model, an engineered construct of LetB, lacking its native transmembrane anchor but chemically tethered at both termini to liposomes, was shown to mediate GPL transport (Cheng et al. 2025). This activity was abolished when either anchor was removed, nonetheless supports a potential role for LetAB in GPL transport. In contrast, no direct biochemical evidence for GPL transport by the Pqi complex has yet been reported.

The Pqi complex is encoded by the *pqiABC* operon, which is induced by paraquat in a SoxS-dependent manner and by starvation via RpoS, suggesting a role in stress adaptation (Koh and Roe 1995, Koh and Roe 1996). PqiA is an IM protein homologous to LetA, although its function remains unclear. PqiB is an MCE domain-containing protein anchored to the IM via transmembrane helices, forming a hexameric assembly of three stacked MCE rings capped by an extended α-helical needle that is proposed to span the periplasm (Ekiert et al. 2017). We have previously shown that PqiB interacts with PqiC, an octameric toroidal lipoprotein localised to the OM, and is required for productive interaction with a GPL bilayer (Cooper et al. 2024).

Despite these insights, the function of the Pqi complex remains unresolved. To date, evidence for its involvement in GPL transport is limited to the co-purification of GPLs with a soluble construct of PqiB (Ekiert et al. 2017) and the combined deletion of *pqiB* and *letB* in a *ΔmlaD* background increasing sensitivity to EDTA, with the overlap in function of the Mla and Pqi systems demonstrated (Nakayama and Zhang-Akiyama 2017). Additionally, Ekiert *et al*. identified OM ruffling in the combined deletion of *mlaE*, *pqiA* and *letA (Ekiert et al. 2017)*. Overall, these results indicate that PqiABC plays a role in maintaining membrane integrity, however the exact function remains elusive.

Here, we develop and structurally resolve a surface based double bilayer Gram-negative envelope mimetic system, complemented by solution-based fluorescence assays, to directly interrogate PqiABC function. We show that PqiABC forms a stable, envelope-spanning complex anchored in both membranes, consistent with a continuous conduit architecture, and that it mediates passive GPL transport, enabling bidirectional lipid exchange between the inner and outer membranes. Together, these results establish PqiABC as a membrane-bridging GPL transport system and provide direct evidence for passive GPL equilibration across the bacterial envelope. Additionally providing a self-assembled didermic envelope biomimetic suitable for molecular-level structural and biophysical studies on inter-membrane transport processes.

## Materials and Methods

### Overexpression of PqiAB, PqiB and PqiC

For protein purification, a full-length construct of PqiC with a C-terminal StrepTag II (named PqiC-Strep) was used throughout (Cooper et al. 2024). An N-terminally Strep-tagged construct of PqiAB was also produced via a Q5 Site directed mutagenesis kit (New England Biolabs, #E0554S) by removal of the C-terminal hexa-histidine tag (His-tag) from the PqiAB construct detailed in (Cooper et al. 2024), followed by insertion of WSHPQFEK at the N-terminus, named PqiAB-Strep (See Supplementary Table S1 for primers used in this study). A construct of PqiB was chemically synthesised (Genscript), consisting of the DNA of PqiB residues 1–546 with a C-terminal His-tag cloned into the pET22b plasmid between the NdeI and XhoI restriction sites. Plasmids were transformed into the *E. coli* C43 (DE3) cell line (Miroux and Walker 1996) and plated on Luria Broth (LB) (Melford) agar, supplemented with 30 µg/mL kanamycin, or 100 μg/mL ampicillin.

Cultures were grown in LB supplemented with the appropriate antibiotic (30 µg/mL kanamycin or 100 μg/mL ampicillin) throughout. 20 mL overnight cultures were used to inoculate 2 L cultures and were grown to an OD_600_ of 0.4 at 37 °C, 180 rpm. For PqiAB overexpression, media was also supplemented with 2.5 mL/L trace metal mix (130 mM Na_2_EDTA.2H_2_O, 80 mM ZnSO_4_.7H_2_O, 180 mM H_3_BO_3_, 40 mM MnCl_2_.4H_2_O, 7 mM CoCl_2_.6H_2_O, 6 mM CuSO_4_.5H_2_O, 0.8 mM (NH_4_)_6_Mo7O_24_.4H_2_O). Cultures were then cooled to 18 °C until cells reached OD_600_ 0.6. Overexpression was induced with the addition of 1 mM isopropyl-β-D-thiogalactopyranoside (IPTG). Protein overexpression was left to proceed overnight at 18 °C, 180 rpm. Cells were harvested via centrifugation (15 min, 5,000 x *g*, 4 °C).

### Purification of PqiAB and PqiC

Cell pellets were resuspended in 20 mM Bis-Tris propane (BTP), 500 mM NaCl, 1 mM EDTA, 0.5 mM tris(2-carboxyethyl)phosphine (TCEP) (pH 8.5) supplemented with 1 c0mplete™ EDTA-free protease inhibitor cocktail tablet (Roche), at a ratio of 25 mL per Litre of overexpression culture. Bacterial cells were lysed via 3-4 passes through an Emulsiflex-C3 cell disruptor (Avestin) at approximately 17,000 psi. Cell debris was removed via centrifugation (30 min, 10,000 x *g*, 4 °C), and membranes harvested via further ultra-centrifugation (1 hr, 100,000 x *g*, 4 °C).

Harvested membranes were resuspended and homogenised in 1 mL per 40 mg of membrane in 20 mM BTP, 150 mM NaCl, 1 mM EDTA, 1 % w/v DDM, 0.5 mM TCEP (pH 8.5). Solubilisation was left to proceed rotating at 4°C for 2 hours. Insoluble material was removed via centrifugation (25 min, 75,000 x *g*, 10°C) and the supernatant subsequently filtered through a 0.45 µm syringe filter. Solubilised protein was then bound to a 5 mL StrepTrap HP column (GE Healthcare), pre-equilibrated in wash buffer (20 mM BTP, 150 mM NaCl, 1 mM EDTA, 0.03 % w/v DDM, 0.5 mM TCEP, pH 8.5). The column was then washed with wash buffer and eluted in 1 mL fractions in wash buffer supplemented with 2.5 mM desthiobiotin. Fractions were analysed by SDS-PAGE and those containing protein of interest were either buffer exchanged using a PD-10 column (Cytiva) into SEC buffer (20 mM BTP, 150 mM NaCl, 0.03 % w/v DDM, 0.5 mM TCEP, pH 8.5) or concentrated using Amicon Ultra 100kDa MWCO centrifugal filters (Millipore) then further purified on a Superose 6 10/300 gel filtration column in SEC buffer.

### PqiAB Amphipol Solubilisation

Amphipol A8-35 (Thermo Fisher) at 50 mg/mL in dH_2_O was mixed with detergent-purified protein at a 4:1 (mg) ratio of Amphipol to protein. The solution was mixed overnight at 4 °C, before excess detergent was removed by Bio-Beads SM-2 resin (Bio-Rad), mixing at 4 °C for 1-2 hrs, removed via a 0.45 µm syringe filter and run on a Superose 6 10/300 column (Cytiva) in 20 mM BTP, 150 mM NaCl, pH 8.5.

### Liposome and Proteoliposome Preparation

Liposomes were produced via the resuspension of dried lipid films to a concentration of 0.1 mg/mL. PqiC-containing proteoliposomes were prepared as detailed in (Cooper et al. 2024) at concentrations between 0.1:0.02-0.04 mg/mL of lipid to PqiC-Strep. Throughout this study proteoliposomes and liposomes were resuspended in 20 mM BTP, 150 mM NaCl (pH 8.5) (supplemented with 1 mM CaCl_2_ throughout NR experiments). The following lipids were used: hydrogen (h) labelled 1-palmitoyl-2-oleoyl-sn-glycero-3-phosphocholine (hPOPC), deuterium (d) labelled Dimyristoyl-d54-sn-glycero-3-phosphocholine (dDMPC) and Dipalmitoyl-d62-sn-glycero-3-phosphocholine (dDPPC).

### Double bilayer assembly

The build-up of a double bilayer system during QCM-D and NR experiments was split up into a series of stages. Surfaces were pre-cleaned via UV/ozone for 10 minutes, immersed in 2% sodium dodecyl sulfate for 30 minutes, rinsed with milliQ water, dried, and UV/ozone treated for a further 10 minutes immediately prior to use. Then PqiC-containing proteoliposomes (consisting of either hPOPC, dDMPC or dDPPC) at a lipid:protein ratio of between 0.1:0.02-0.04 mg/mL were introduced by flow onto the silicon surface then ruptured via osmotic shock to create a proteolipid bilayer on the silicon surface with PqiC exposed on the external facing leaflet. Next, the addition of 0.01-0.04 mg/ml PqiAB (stabilised with amphipol A8-35) was added to the system leading to binding to the surface exposed PqiC and PqiABC complex formation. Finally, a second hPOPC bilayer was generated through the addition and subsequent osmotic shock of 0.1 mg/mL hPOPC liposomes. Throughout these experiments samples were resuspended in 20 mM BTP, 150 mM NaCl (pH 8.5) (supplemented with 1 mM CaCl_2_ throughout NR experiments).

**Quartz crystal microbalance with dissipation monitoring (QCM-D)** is a surface-sensitive method that exploits the piezoelectric properties of quartz to track interfacial processes in real time. The technique records shifts in the resonance frequency of a quartz crystal sensor; decreases in frequency (Δf) scale inversely with the mass adsorbed at the surface, as described by the Sauerbrey relationship:

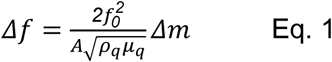

Where *f_0_* represents the resonant frequency, *A* is the piezoelectrically excited area of the crystal and *ρ_q_* and *μ_q_* correspond to the density and shear modulus of the quartz crystal, respectively.

Measurements were carried out using a Q-Sense explorer (Biolin Scientific). Experiments were performed at 25 °C unless otherwise stated with a constant flow rate of 0.1 mL/min. Both frequency and dissipation shifts (Δf and ΔD) were recorded across multiple overtones (n = 3, 5, 7, 9, 11, and 13). For clarity, only n = 3, 5 and 7 were plotted. Data acquisition began in H_2_O, followed by exchange into buffer containing 20 mM BTP and 150 mM NaCl (pH 8.5) and subsequent bilayer assembly as detailed above.

### Neutron reflectometry

Specular NR measurements were performed on the white beam OFFSPEC (Campana et al. 2026) reflectometer at the ISIS Neutron and Muon Source (Oxfordshire, UK) to structurally examine the assembly and distribution of protein and GPL components across Pqi complex separated double membrane samples. Solid-liquid (S-L) flow cells containing a 80 mm × 50 mm × 15 mm silicon substrate with a single 50 mm × 80 mm face polished to 3 Å rms roughness in contact with a solution containing trough (Clifton et al. 2019) were placed on the instrument sample stage and aligned in terms of height and angle to the incoming neutron beam. NR measurements were obtained over a Q_z_ range ∼0.01 to ∼0.3 Å^-1^ using OFFSPEC’s instrument wavelength band of 1 to 14 Å (λ) and angles (θ) of 0.6°, 1.2° and 2.3° of the neutron beam relative to the silicon-water interface, where:

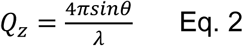

The solution Hydrogen/Deuterium isotopic contrast within the S-L flow cells was changed using an instrument dedicated liquid chromatography pump (LC-4000, Jasco, Japan) at a rate of 1 mL/min and the temperature of the S-L flow cells was controlled using an instrument dedicated circulating water-bath (FP50, Julabo, Germany).

Initially the bare silicon-solution interface was measured under two buffer (20 mM BTP, 150 mM NaCl, pH 8.5) solution isotopic contrast conditions (D_2_O and H_2_O) thereafter the membrane-protein samples were measured under three buffer solution isotopic contrast conditions (D_2_O, H_2_O and Protein-matched water (PrMW, 42% D_2_O)). The neutron spectra obtained at each reflecting angle were converted to reflectivity profiles by dividing by the non-reflected neutron transmission spectra and then stitching of the individual angular data sets into a single reflectivity profile.

**NR data analysis** for the Pqi-containing double bilayer samples and their step-wise assembly was conducted using the RasCal software (Hughes 2019) which uses Abeles layer models (Abelès 1948) to describe the neutron scattering length density (nSLD) profiles across bulk interfaces. Rascal’s custom model option was used to define models which resolved the sample structural parameters (macromolecular component volumes, sizes and distributions) by using these to calculate model reflectivity data sets and nSLD profiles which were fitted to the experiment reflectivity data. Stages 1-3 of the assembly process were fitted independently using the three solution isotopic contrast data sets obtained for each experimental stage. Data sets from all stages were simultaneously fitted with the NR data from the bare silicon surface. Model-to-data fits were initially optimised using a simplex algorithm with Chi^2^ used as a goodness of fit guide followed by RasCal’s Monte Carlo Markov Chain Bayesian inference routines to estimate structural ambiguities.

NR data was fitted using density profiles derived from the published PDB structure of PqiC (see Supplementary Figure S1) and an alphafold model of PqiABC (Jumper et al. 2021)(see Supplementary Figure S2) to produce the Pqi scattering length density profiles above the proximal bilayer. To incorporate the structures of PqiC and PqiABC into the NR data fitting, 1-dimensional density profiles of the proteins were generated along the proteins minor axis for PqiC and major axis for PqiABC in 5 Å bins (see Supplementary Figure S1 and S2) and normalised to the region of highest chain density across the distributions. These 1-D density profiles for the protein structures were then converted into a series of layers in a RasCal custom model with the protein density profile scaled relative to a fitted coverage parameter for the proteins content on the proximal bilayer surface.

The position of the distal bilayer GPLs (d and h-lipid) on the PqiABC complex was fitted as the fraction of the total length of the Pqi complex where the component distributions began. The fitted density of these components was then added to these regions of the protein structure until the end-position minus the start-position along the PqiABC structure was equal to the fitted thickness of the distal bilayer. The minor distributions of lipid found following Pqi-complex assembly was fitted as a single nSLD value (Arunmanee et al. 2018) whereas the distal bilayer structure following hPOPC addition was fitted as a lipid head-lipid tail-lipid head distribution (Clifton et al. 2019) with the nSLD value of the tails calculated from the fitted d/h-lipid mixture in this region of the interfacial structure (see Supplementary Table S2). A similar, but independent mixing, parameter was used to fit the nSLD of the inner bilayer. In this case, the d/h-lipid content of the two leaflets of the tails was fitted independently to allow for substrate induced asymmetry (Wacklin 2011). The final fits of the interfacial structure were conducted using RasCals resampling function which re-parameterises the interfacial layer structure as a series of interrelated thin (1 Å) un-roughened slabs to prevent problems with large interfacial layer roughness relative to layer thickness (John et al. 2021).

Once satisfactory fits were obtained, error analysis was performed on the model-to-data fits to assess the ambiguity of the resolved interfacial structures. This was achieved using Rascals in-built Bayesian error estimation routines. Bayesian inference was undertaken using MCMCStat (https://mjlaine.github.io/mcmcstat) Delayed-Rejection Adaptive Metropolis algorithms (DRAM) (Haario et al. 2006) Monte-Carlo-Markov Chain (MCMC) routines to refit the data using a user defined number of links. To fit data using this approach, the likelihood function is defined in terms of the Chi-squared goodness of fit criteria, as shown previously (Sivia and Webster 1998). The parameter uncertainties were then determined from the MCMC derived posterior distributions as the shortest percentile confidence interval (Sivia and Skilling 2006) from each (in this case 65%). The uncertainties within this interval for the reflectivity fits and SLD profiles were generated by randomly sampling (in this case 1000 samples) from the Markov chains, calculating reflectivity’s and SLD’s for each set of samples, and taking the relevant percentile across all the resampled reflectivity or SLD curves at each point in Q_z_ or distance to represent the uncertainties on the fits. The median of the reflectivity and SLD uncertainties are shown as a darker line in figures.

Volume fraction profiles detailing the distribution of components across the solid/liquid interface were produced using a bespoke script. Parameter distributions from the Bayesian error estimation process and the relationship between the fitted parameters and interfacial structure, as defined in the RasCal custom model, were used to determine the distribution of each structural component across the solid-liquid interface. The volume fraction of an individual component was calculated in 1 Å increments across the surface (the silicon/silicon dioxide interface set as zero). The median, lower, and upper 65% confidence interval bounds of each component distribution were determined for every 1 Å segment using the MCMC error estimation results or derived parameters; these confidence intervals were then used to produce line width error regions above and below the median values. The water distribution was calculated as the remaining unoccupied volume for each 1 Å slice and summed across the interface with the appropriate error propagation.

### Assessment of Interbilayer Mixing Using Time NR

During temperature/lipid phase change induced proximal and distal bilayer mixing studies on a dDPPC-PqiABC-hPOPC sample (Figure 4) the depth of the 1st Kiessig fringe minima (∼0.029 Å^-1^) in the H_2_O solution contrast was used as a diagnostic of the evolution of this process as it was the most prominent feature change during NR analysis of lipid transfer between the bilayers (see Figure 4A). The depth of this minima in the data sets collected at 20℃ and 42℃ were defined as 0 and 100% of the total mixing process respectively. The change in data during the heating of the sample from 30-to-34℃ was chosen to probe the temporal relationship between bilayer mixing and sample temperature as the biggest change in NR was found during this heating stage (∼47 to ∼90% of the complete exchange process). NR data collected during the heating process was reduced into a series of the 120 second time sliced data sets and the depth of the minima determined as a percentage of the total mixing process, with appropriate error propagation. The time dependent change in the bilayer mixing process during heating was then compared to the temperature read back value from the sample heating water bath temperature which was collected concurrently with the NR data.

### FRET-based GPL transport assays

GPL mixtures were prepared from chloroform stocks and dried under a stream of air in glass vials and stored at −20 °C. For FRET-based studies, headgroup labelled 1,2-dioleoyl-sn-glycero-3-phosphoethanolamine-N-(7-nitro-2-1,3-benzoxadiazol-4-yl) (NBD-PE) and 1,2-dipalmitoyl-sn-glycero-3-phosphoethanolamine-N-(lissamine rhodamine B sulfonyl) (Rhod-PE), and POPC were used. Dried GPL mixes were resuspended to a final concentration of 250 µM in the assay buffer (20 mM BTP, 150 mM NaCl, pH 8.5).

A master mix was prepared at a volume ratio of 11.5:1:1 (assay buffer:donor liposomes:acceptor liposomes), and 135 µL was dispensed into each well of a black 96-well microplate. Donor liposomes comprised 97:1.5:1.5 mol% POPC:NBD-PE:Rhod-PE, while acceptor liposomes consisted of 100 mol% POPC. Baseline fluorescence was recorded until a stable signal was established. Protein stock (15 µL) or an equivalent volume of SEC buffer was then added to each well, ensuring that the final detergent concentration remained below the critical micelle concentration (CMC) of DDM. A fully mixed positive control was generated by adding 15 µL of 10% (w/v) DDM in assay buffer in place of protein or SEC buffer, resulting in complete liposome solubilisation and maximal fluorophore mixing. Plates were immediately returned to the plate reader, and fluorescence was monitored over time at 25 °C.

Unless otherwise stated, protein stocks were standardised to a concentration of 1 µM in SEC buffer before addition to the assay. Where indicated, Pqi(A)B and PqiC were incubated together overnight at 4°C before addition to the assay to allow complex formation. Gain-normalised fluorescence measurements were acquired using a TECAN Infinite 200 plate reader with excitation at 460 nm and emission at 535 nm. All measurements were performed in technical triplicate.

Data was normalised as below:

Dequenching assay normalisation:

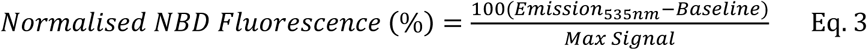

Where the max signal is the average 10 % DDM sample over time.

The mean and standard error of the mean over time were plotted using GraphPad (Prism).

### *In vivo* phenotypic complementation assays

Constructs of PqiAB and PqiABC in pET26b plasmids and PqiC in pET22b, each with a C-terminal His-tag, in addition to the PqiB construct detailed above, were used as detailed by (Cooper et al. 2024) for complementation assays. Plasmids were transformed into chemically component *E. coli* K-12 BW25113 cell lines (Baba et al. 2006) for phenotypic complementation studies, containing a knockout of genomic *pqiABC* (Cooper et al. 2024). Single colonies were picked to inoculate 5 mL overnight cultures in LB supplemented with the appropriate antibiotic (30 µg/mL kanamycin or 100 µg/mL ampicillin). Serial dilutions of the overnight cultures were set up from OD_600_ 10^0^ to OD_600_ 10^-7^ in LB. 2.5 µL of the serially diluted cultures were spotted onto LB agar plates supplemented with 0.25 % (w/v) lauryl sulfobetaine (LSB) and dried at room temperature for 3 hours before overnight incubation at 37°C. Plates were imaged using ChemiDoc XRS+ (Bio-Rad).

### Western blots to detect protein expression in complementation assays

Whole cell lysate samples were produced to assess leaky protein expression in phenotypic complementation assays. 1 mL of overnight culture was pelleted by centrifugation 4,000 x g, 10 min, and the cell pellet was resuspended in 100 µL 50 mM Tris (pH 8), 150 mM NaCl, 2 % Triton 100, and incubated on ice for 30 min. Cell debris was pelleted at 10,000 x g, 10 min and 60 µL supernatant mixed with 20 µL 4x Laemmli Sample Buffer.

Samples were loaded onto SurePAGE™, Bis-Tris, 4–12% polyacrylamide gels (GenScript) alongside a peqGOLD Protein Marker IV (VWR). Gels were run in MES SDS running buffer (GenScript) at a constant 165 V until complete. Samples were then transferred to a Poly(vinylidene fluoride) membrane using a Trans-blot Turbo pack (Bio-Rad), 1.3 A, 25 V, 7 min. Membranes were blocked in 5 % bovine serum albumin (BSA) in 1 x TBST (20 mM Tris, 150 mM NaCl, 0.1 % w/v Tween 20) for 60 min shaking at room temperature. Membranes were then incubated in a 6x His monoclonal (anti-His) antibody (Takara Bio), diluted 1:2500 in 5 % (w/v) BSA solution in 1 x TBST, mixing for 60 min at room temperature. Membranes were washed 3 x 10 min in 1 x TBST, before the addition of anti-mouse HRP-conjugated Secondary antibody (ThermoFisher) in 1 x TBST. Membranes were then shaken at room temperature for 60 min, followed by 3 x 10 min washes in 1 x TBST. Blots were treated with Amersham™ ECL Western Blotting Detection Reagent (Cytiva) and imaged using chemiluminescence on Amersham Imager 680 (GE healthcare).

## Results and Discussion

### PqiABC forms an envelope spanning complex, anchoring to both the inner and outer membranes

To investigate the structure of PqiABC and its interactions with the inner and outer membranes, we combined quartz crystal microbalance with dissipation monitoring (QCM-D) and neutron reflectometry (NR). Using these complementary techniques, we reconstructed PqiABC from its individual components (PqiAB and PqiC) within a silicon-supported, double bilayer mimetic. QCM-D enabled real-time monitoring of bilayer formation through mass changes at the sensor surface, while NR, exploiting isotopic contrast variation, resolved the distributions of protein, lipid, and solution across the interfacial assembly. Together, these approaches allowed us to define the assembly and organisation of the complex, while providing a solution-phase double bilayer platform for interrogating Pqi function under physiologically relevant conditions.

We constructed the system in a stepwise manner, assessing the interfacial structure at each stage using NR. Figure 1 illustrates the overall assembly process including QCM-D traces, NR data, model fits, the resulting scattering length density (SLD) and component volume fraction profiles. The assembly was carried out in three stages (Figure 1A): (1) formation of a protiated 1-Palmitoyl-2-oleoyl-sn-glycero-3-phosphocholine (h-POPC) planar supported lipid bilayer (SLB) with surface-bound PqiC directly on a silicon support; (2) binding of PqiAB, solubilised in A8-35 amphipol, to PqiC on the hPOPC membrane; and (3) a second addition of hPOPC, causing the formation of a high coverage second lipid bilayer. POPC was selected for its ability to form uniform, defect-free planar bilayers and its low gel–fluid transition temperature, ensuring a fully fluid membrane at room temperature. While not compositionally representative of bacterial membranes, these properties provide a stable and well-defined platform that captures key physical characteristics of the liquid-crystalline state. Consequently, POPC has become one of the most widely used model phospholipids for investigating membrane-associated processes in planar bilayer systems (Puu and Gustafson 1997, Castellana and Cremer 2006, Andersson and Koper 2016).

**Figure 1.**
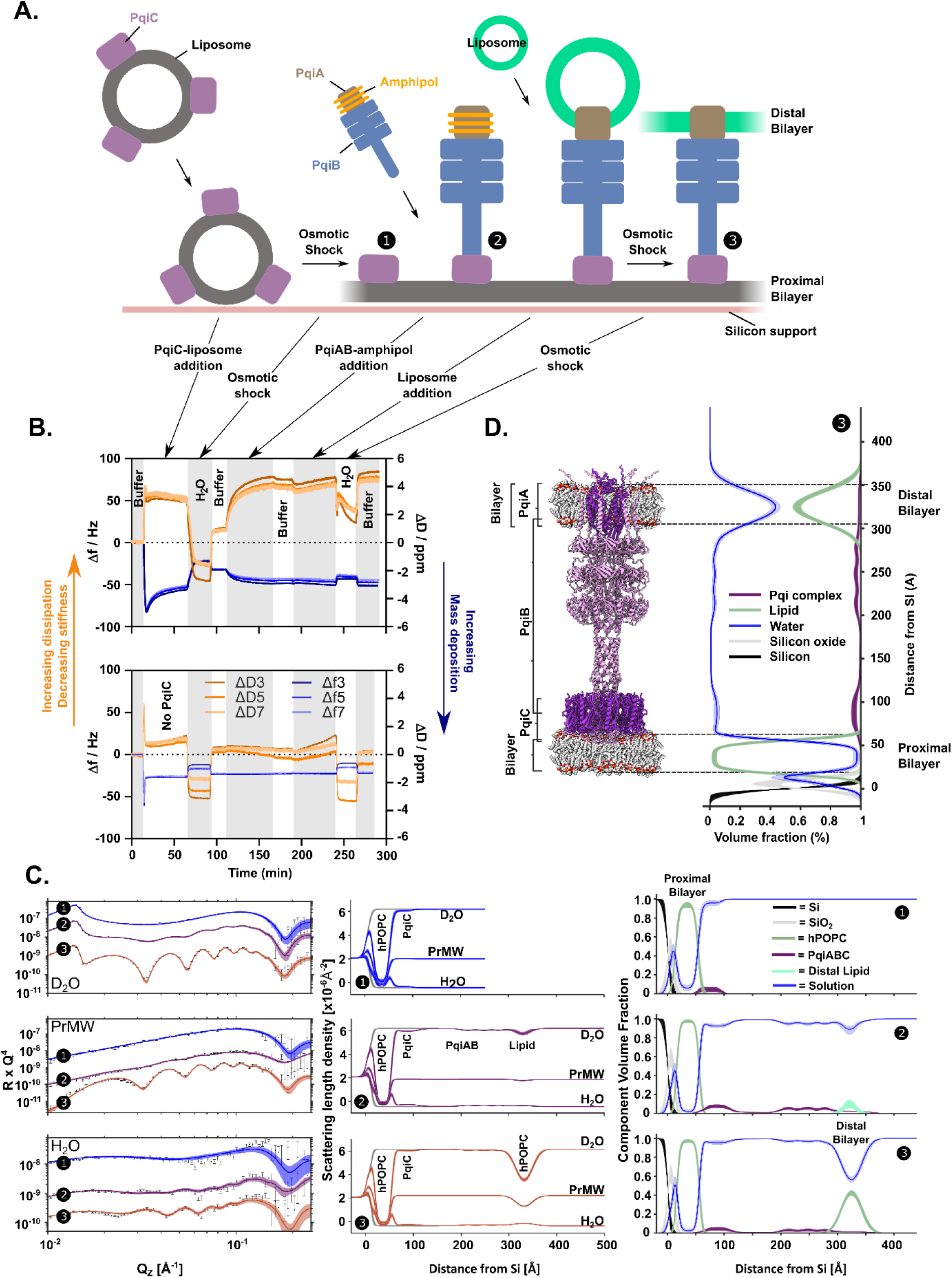
PqiABC forms a transenvelope complex linking the inner and outer membranes. **(A)** Schematic of the silicon-supported GPL double bilayer deposition process showing the three stages of deposition: ❶ PqiC-hPOPC planar bilayer deposition, ❷ PqiAB-amphipol addition, ❸ hPOPC addition to form a distal bilayer. **(B)** corresponding QCM-D of the deposition process, highlighting PqiAB deposition requires the presence of PqiC. **(C)** NR profiles (left, error bars) and model data fits (left, lines), the scattering length density (SLD) profiles (middle) and component volume fraction (CVF) *vs.* distance profiles (right) these fits describe during the assembly process described in (A). **(D)** Component volume fraction vs distance profile of ❸ mapped to the alphafold model of PqiABC. The line widths in the NR model data fits depict the range of acceptable fits within the 65% confidence interval range of the posterior distributions of the fitted model parameters after MCMC error analysis. Ambiguity in the resolved surface structure is shown as line widths in the SLD and CVF profiles which are derived from the 65% confidence interval of the parameter posterior distributions.

Stage 1 QCM-D measurements, consistent with our previous work (Cooper et al. 2024), indicated that the addition of PqiC-hPOPC proteoliposomes (1:2.5 w/w) led to a decrease in resonant frequency with a maximum decrease of −80 Hz observed with a concomitant increase in dissipation (∼3 ppm) (Figure 1B). This response is characteristic of the adsorption of intact, viscoelastic lipid vesicles onto the sensor surface (Cho et al. 2010). A subsequent partial rupture of the liposomes was noted, evidenced by the increase in frequency observed. To complete the vesicle rupture process, an osmotic shock was performed, with subsequent return to the buffer. Overall, this resulted in a frequency decrease of ∼33 Hz for the GPL deposition process. This is considerably larger than expected for deposition of a complete supported POPC bilayer alone (typically Δf ≈ −25 to −27 Hz in QCM-D), indicating the presence of additional adsorbed material beyond a single planar bilayer (Cho et al. 2010), and is consistent with the additional PqiC within the system (Cooper et al. 2024). To confirm the interfacial structure generated by the deposition process, NR analysis was performed, revealing the formation of a PqiC-containing planar bilayer with ∼97% surface coverage with a ∼5% coverage of the PqiC octamer asymmetrically located on the bilayer’s, solution facing, outer leaflet (Figure 1C-❶ & Table 1). From here on in, we refer to this membrane as the proximal bilayer.

**Table 1.** A comparison of the coverages and composition of proximal and distal bilayer hPOPC GPL coverages and their separation by the PqiABC complex.

| Region | Parameter | Percentage coverage (%) / Distance (Å) |
| --- | --- | --- |
| Proximal Bilayer Coverage | Lipid | 97% (95–99) |
|  | Solution | 3% (1–5) |
| Inter-Membrane Region | ① - PqiC (membrane surface) | 5% (2–8) |
|  | ② and ③ - PqiABC (membrane surface) | 6% (4–8) |
|  | Inter-bilayer Distance (Å) | 255 Å (250–260) |
| Distal Bilayer Coverage | Lipid | 59% (52–68) |
|  | PqiABC (distal bilayer region) | 1% (1–1) |
|  | Solution | 41% (32–48) |

Following on from stage one, PqiAB solubilised in A8-35 amphipol was introduced to the system. QCM-D showed clear evidence of binding to the planar surface, with a decrease of ∼15 Hz in resonant frequency observed. This binding event was mediated by the presence of PqiC, as in its absence, no binding was observed (Figure 1B). This is consistent with our previous data using PqiAB-containing proteoliposomes (Cooper et al. 2024). Such results suggested stable formation of the full PqiABC complex. NR analysis confirmed this, showing PqiABC complex formation on the proximal bilayer surface (Figure 1C-❷), with analysis suggesting the protein complex was orientated perpendicular to the SLB surface i.e. along the surface normal. The coverage of PqiABC, i.e. the area its chains occupied on the SLB surface, was found to be ∼6%, consistent with PqiC levels on the surface (Table 1). This was based on the maximum density of the complex, with the volume fraction of protein in the complex varying with distance along the solid-liquid interface (Supplementary Figure S2). As the total coverage of the Pqi complex on the proximal bilayer surface was determined from protein chain density by NR, the actual coverage can be estimated to be nearly double the chain values when protein associated water is accounted for (Clifton et al. 2016).

A distribution of hydrogenous material, distinct from the protein density, was clearly resolved within the hydrophobic belt region of PqiAB following assembly of the PqiABC complex. The SLD of this material was consistent with either amphipol or GPL (Figure 1C-❷ and Supplementary Table S2). While this density was initially assigned to amphipol, comparison with samples in which the proximal bilayer GPLs were maintained in the gel phase (see Supplementary, Section 2) revealed that this density was not present. These observations indicate that the material is more likely to arise from GPLs already present within the system, originating from the proximal bilayer, or native GPLs co-purified with PqiAB, or both, which form low density distal bilayer when the sample is in the fluid phase.

Next, we examined if the addition of hPOPC vesicles to the sample followed by osmotic shock would cause the formation of a higher coverage lipid bilayer around the PqiAB’s hydrophobic belt. QCM-D results showed that addition of hPOPC minimally impacted the resonant frequency but led to a larger change in dissipation, suggesting that, rather than the binding of vesicles, an exchange of material had occurred changing the viscoelastic properties (Figure 1B). Subsequent osmotic shock had minimal impact on both frequency and dissipation further confirming that exchange was likely rather than vesicle binding. NR confirmed displacement and the formation of a second membrane, hereafter referred to as the distal bilayer (Figure 1C-❸ & 1D), located ∼265 Å from the proximal membrane. This distance is consistent with the expected size of the *E. coli* periplasmic space and with the overall length of the Pqi complex, which spans ∼320 Å across the reconstituted envelope. The surface coverage of the distal bilayer was ∼60% (Table 1), complete coverage was not expected given the relatively low (∼6%) surface coverage of PqiABC (Table 1) which provides the supporting scaffold around which this bilayer forms.

Previous work from our laboratories developing free-floating bilayer systems demonstrated that bilayer fluctuation amplitudes increase markedly with distance from the underlying substrate (John et al. 2021, Hall et al. 2024). For example, in the absence of a protein scaffold, a free-floating bilayer positioned approximately 200 Å from the substrate exhibited a roughness of ∼63 Å, reflecting substantial thermal fluctuations and membrane undulations (John et al. 2021). In contrast, the use of PqiAB as a scaffold to anchor the distal bilayer appears to restrict the fluctuational freedom of the fluid-phase membrane, exhibiting only a modest z-directional broadening, corresponding to a roughness of approximately ∼9 Å at 260 Å from the proximal bilayer and >300 Å from the support. The remaining roughness most likely reflects a combination of residual thermal fluctuations and undulations of the distal GPL bilayer (John et al. 2021), together with minor sample heterogeneity across the 30 × 60 mm NR sample surface. This sevenfold reduction in roughness, despite the larger inter-bilayer spacing, strongly suggests that PqiAB mechanically constrains the distal membrane, confining its motion within the interfacial architecture.

### PqiABC allows passive mixing of glycerophospholipids between the proximal and distal membranes

To investigate whether PqiABC enables passive GPL mixing between bilayers, we reconstructed the system, this time exploiting the isotopic contrast between protiated and deuterated lipids. Deuterated 1,2-dimyristoylphosphocholine (dDMPC) was incorporated into the proximal bilayer, while protiated POPC (hPOPC) was used for the distal bilayer (Figure 2; Table 2). dDMPC was initially selected due to its availability and cost-effectiveness. As NR is sensitive to hydrogen isotopes, this labelling strategy allowed quantification of GPL distribution across the two bilayers while also enhancing the contrast between the protiated protein and the deuterated membrane.

**Figure 2.**
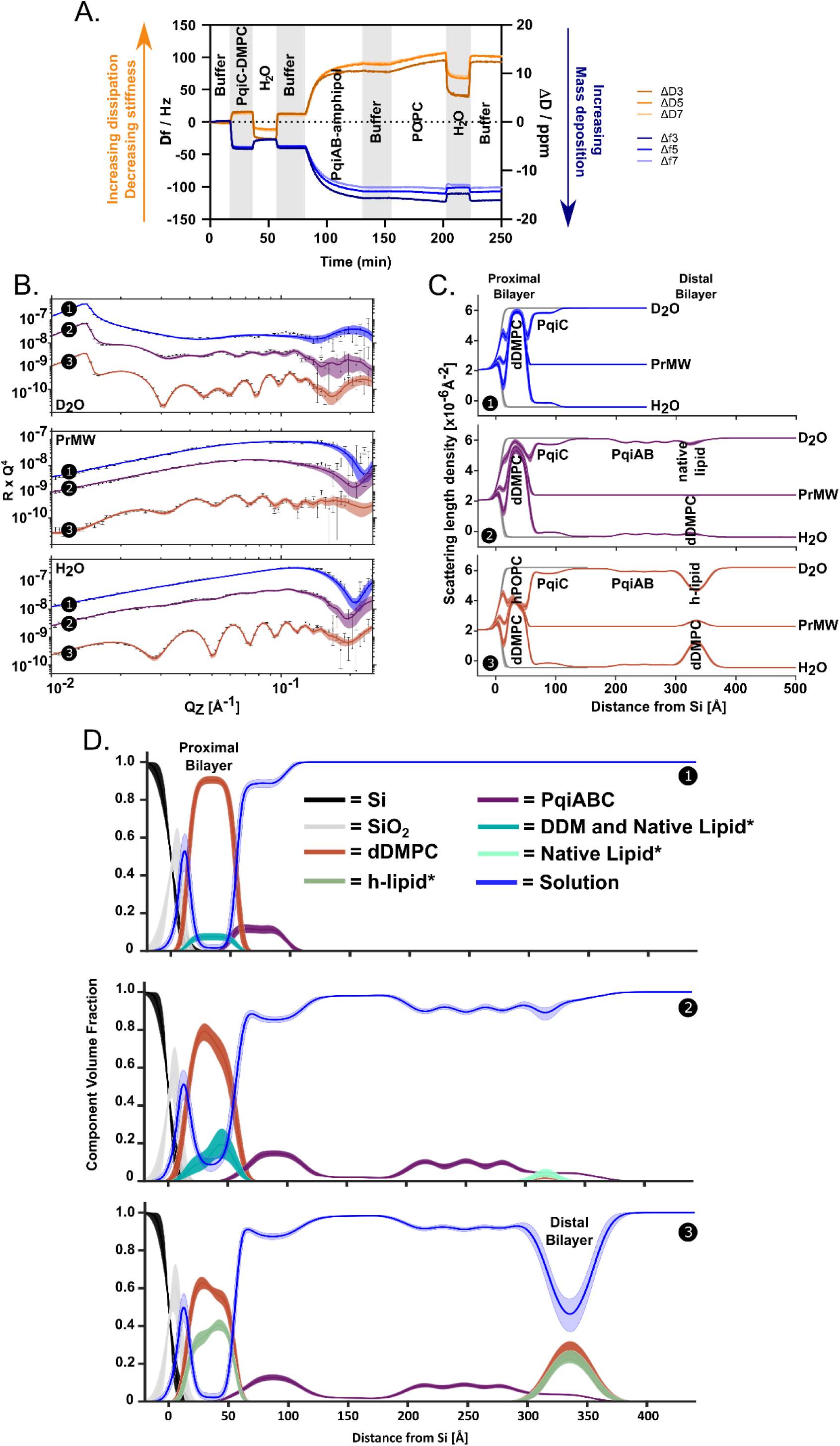
Differential hydrogen/deuterium labelling of proximal and distal bilayer GPLs reveals mixing between the Pqi-separated bilayers. QCM-D of the deposition process (A), consistent with the schematic shown in Figure 1A with dDMPC used for proximal bilayer deposition and hPOPC used for distal bilayer deposition. NR profiles (B, error bars) and model data fits (B, lines) and the scattering length density profiles these fits describe (C) obtained during the assembly of a double GPL bilayer spanned by PqiABC. Initially a dDMPC proximal bilayer with outer surface of PqiC was deposited onto the Si surface (NR fits and SLD profiles are shown in blue and ❶). Onto this amphipol-stabilised PqiAB proteins were added from solution where they bound to PqiC (purple and ❷). Finally, hPOPC was deposited from a vesicle suspension where it displaced amphipol from the PqiAB’s membrane spanning region forming the second distal membrane ∼260 Å away from the proximal bilayer (red and ❸). Fitting derived component volume fraction vs. distance profiles for the assembly of the Pqi separated double bilayer sample are shown (D). The line widths in the NR model data fits depict the range of acceptable fits within the 65% confidence interval range of the posterior distributions of the fitted model parameters after MCMC error analysis. Ambiguity in the resolved surface structure is shown as line widths in the SLD and volume fraction profiles which are derived from the 65% confidence interval of the parameter posterior distributions. *Denotes contributions to the interfacial structure which are inferred from the fitted nSLD values.

**Table 2.** NR-fitting derived structural parameters obtained for the initial stages (1 and 2) of the assembly of a PqiABC spanning double bilayer sample with a proximal bilayer deposited as dDMPC and a distal bilayer deposited as hPOPC.

| Stage | Proximal bilayer thickness (Å) | Proximal bilayer composition (%) | Pqi coverage (%) | Bilayer-Amphipol Distance (Å) | Distal bilayer composition (%) |
| --- | --- | --- | --- | --- | --- |
| 1 | 42(40-44) | dDMPC: 90% (88-92)<br>Contaminant (likely DDM): 7% (6-8)<br>Solution: 2% (0-2) | PqiC: 13% (11-16) | N/A | N/A |
| 2 | 47(44-48) | dDMPC: 76% (69-82)<br>Contaminant (likely DDM and native lipids): 16% (9-23)<br>Solution: 9% (5-14) | PqiABC: 18% (16-20) | 270 | native Lipid 6% (3, 10)<br>dDMPC 4% (1, 11)) |

Bilayer assembly was performed above the gel-to-liquid crystalline phase transition temperatures of both GPLs (dDMPC, 21 °C; hPOPC, −3 °C) to ensure that each bilayer remained in the fluid phase, thereby maximising deposition efficiency. QCM-D indicated that deposition proceeded similarly to the hPOPC:hPOPC (proximal:distal) system (Figure 2A vs. Figure 1B), but with increased incorporation of PqiC (∼40 Hz) and PqiAB (∼77 Hz). This trend was corroborated by NR, which showed ∼13% surface coverage of PqiC compared to ∼5% in the previous system (Figure 2B–D; Table 2).

Interestingly, during proximal bilayer deposition (stage one), a minor (∼10%) protiated contribution was detected within the acyl chain region despite the use of deuterated GPLs (Figure 2C-❶); this feature was likely present but not observable in the original fully protiated system (Figure 1) due to the absence of hydrogen–deuterium contrast. Based on the fitted neutron scattering length density, this signal is consistent with residual DDM carried over from PqiC stabilisation during proteoliposome preparation, where it can intercalate into GPL vesicles.

To test this, proximal bilayer deposition was repeated using a lower PqiC:dDMPC ratio (1:5 rather than 1:2.5, w/w). This resulted in reduced PqiC surface coverage and a corresponding decrease in the protiated contribution (∼4%), consistent with reduced detergent carryover during vesicle preparation (Supplementary Figure S3).

Upon binding of PqiAB–amphipol to PqiC in the proximal membrane, NR further revealed an increase in bilayer hydration (∼8%), consistent with partial insertion of PqiAB and the transfer of some proximal bilayer dDMPC to the low coverage mixed d and h-GPL distal bilayer which appeared after full Pqi complex assembly (Figure 2D-❷).

Across independent experiments (Figures 2, 3 and Supplementary Figure S4), we consistently observed an additional increase in protiated material within the proximal bilayer on PqiAB-amphipol addition, likely reflecting incorporation of co-purified, protiated GPLs associated with the PqiB needle, consistent with previous observations of GPL co-purification with PqiB (Ekiert et al. 2017). However, a contribution from amphipol cannot be excluded, although NR analysis indicates that any amphipol present is likely to constitute only a minor fraction of the observed density (Supplementary, Section 2). Collectively, these data suggest that the hydrogenous material within the proximal bilayer predominantly comprises residual detergent and protein-associated GPLs, with at most a minor contribution from amphipol.

Following the addition of hPOPC vesicles to increase distal bilayer coverage, NR measurements of the completed system revealed that the two bilayers had converged to remarkably similar GPL compositions, despite being separated by ∼265 Å. The solution-facing leaflet of the proximal bilayer (used to minimise support-induced artefacts) comprised 58% dDMPC and 42% protiated material, while the distal bilayer contained 52% dDMPC and 48% protiated material (Figure 2; Table 3). Because hPOPC, DDM and amphipol have similar neutron scattering properties, the protiated fraction cannot be uniquely resolved. It is therefore expected to comprise predominantly hPOPC, together with smaller contributions from residual DDM, amphipol and protiated GPLs co-purified with PqiAB.

**Table 3.** A comparison of the coverages and composition of proximal and distal bilayer lipid coverages and their separation by the PqiABC complex.

| Region | Parameter | Value |
| --- | --- | --- |
| Proximal Bilayer | Lipid Coverage | 98% (96–99) |
|  | ↳ Composition (inner leaflet) | dDMPC: 65% (61–69)<br>hLipid*: 34% (30–38) |
|  | ↳ Composition (outer leaflet) | dDMPC: 58% (55–61)<br>hLipid*: 42% (39–45) |
|  | Solution | 2% (1–4) |
| Inter-Membrane Region | PqiABC Coverage | 16% (14–18) |
|  | Inter-bilayer Distance (Å) | 265 (260–270) |
| Distal Bilayer | Lipid Coverage | 59% (47–72) |
|  | ↳ Composition (bilayer) | dDMPC: 53% (50–56)<br>hLipid*: 47% (44–50) |

The near-equilibration observed in the absence of an energy source is highly supportive of PqiABC facilitating passive, bidirectional GPL transport between the two bilayers. In the absence of such transport, hPOPC would be expected to remain largely confined to the distal bilayer, with only limited incorporation into the proximal membrane through direct vesicle– bilayer interactions. Such interactions are likely to be further restricted by the ∼18% surface coverage of PqiABC protruding into solution (Table 2), which sterically limits access of vesicles to the underlying proximal bilayer. Together, these observations strongly support a protein-mediated mechanism for GPL exchange.

A second, more subtle alternative mechanism is spontaneous lipid exchange via monomer desorption and transfer between membranes. Phospholipids have been shown to undergo spontaneous transfer between lipid vesicles in the absence of lipid transfer proteins, with transfer rates depending strongly on lipid chemistry, particularly acyl chain length (Richens et al. 2017). However, several features of our experimental system make this explanation unlikely. First, the GPLs used were zwitterionic (POPC and DMPC), whereas spontaneous transfer has generally been reported to be more favourable for specific charged phospholipids or under conditions that promote lipid desorption. Second, both POPC and DMPC possess relatively long acyl chains, which substantially reduce spontaneous monomer exchange because of their low aqueous solubility. Finally, the experiments were performed at relatively low temperatures (25 °C), further suppressing lipid desorption and transfer kinetics. Collectively, these considerations suggest that spontaneous lipid mixing is unlikely to account for the observed convergence in bilayer composition.

### Differential lipid phase transition temperatures between proximal and distal membranes allows control of transport

To further confirm that bilayer mixing was mediated by PqiABC, rather than arising from liposome mixing during deposition or spontaneous lipid exchange, we exploited differences in gel-to-liquid crystalline phase transition temperatures between the proximal and distal bilayers to create an activity switch, enabling structural interrogation of PqiABC-mediated GPL transport.

Previously, GPL mixing was observed at 25 °C when both the proximal (dDMPC) and distal (hPOPC) bilayers were in the fluid phase (Figure 2). To decouple this behaviour, the system was reconstructed using the deuterated long-chain (C16) saturated GPL 1,2-dipalmitoylphosphatidylcholine (dDPPC) in the proximal membrane, while retaining hPOPC in the distal membrane. Owing to its elevated phase transition temperature (∼37 °C; pre-transition ∼29 °C (Bryant et al. 2019)) we hypothesised that dDPPC would enable the proximal bilayer to remain in the gel phase under conditions where hPOPC in the distal bilayer remains fluid.

Proximal bilayer deposition (PqiC–dDPPC) and deposition of PqiAB–amphipol was performed at 42 °C, above the main phase transition temperature of dDPPC, to ensure efficient deposition of the bilayer and Pqi-complex formation. Subsequently deposition of hPOPC to form the h-lipid distal bilayer was carried out at 20 °C, below both the pre-transition and main transition temperatures of dDPPC (Figure 3A). Under these conditions, the proximal bilayer should remain in the gel phase, preserving phase asymmetry while allowing efficient deposition of the fluid hPOPC to the distal bilayer.

**Figure 3.**
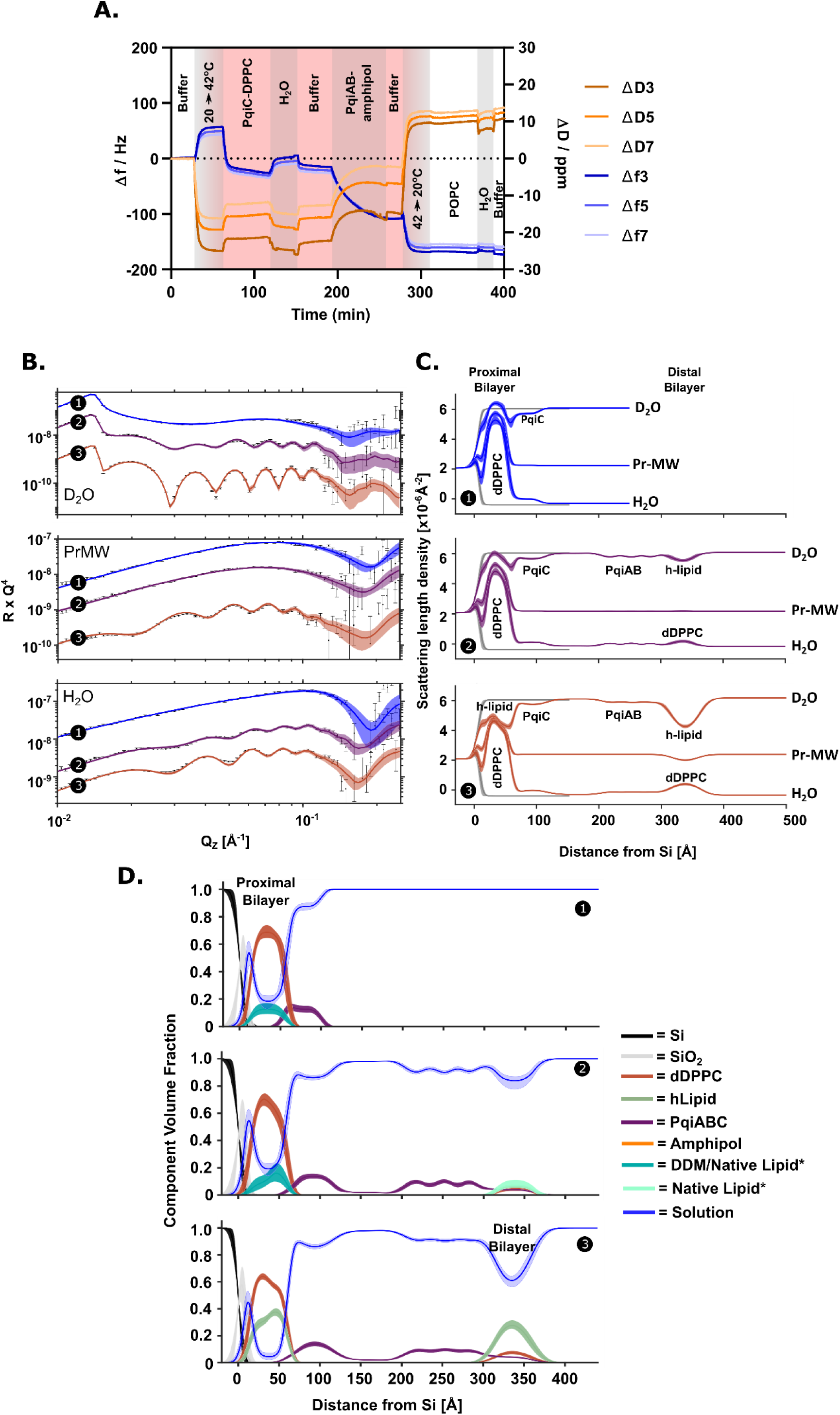
Temperature-driven asymmetric double bilayer deposition using GPLs with distinct phase transition temperatures. **(A)** QCM-D analysis of the deposition process. Temperature was elevated to 42 °C to facilitate deposition of PqiC–DPPC proteoliposomes (red), then decreased to 20 °C for subsequent steps. Deposition was performed as described in Figure 1A. **(B)** NR profiles (error bars) and model data fits (lines) and the scattering length density profiles these fits describe **(C)** obtained during the assembly of a PqiABC separated double membrane sample system with a proximal bilayer deposited as dDPPC and a distal bilayer deposited as hPOPC. Initially a dDPPC SLB with surface PqiC was deposited onto the Si surface (NR fits and SLD profiles are shown in blue and ❶). Onto this PqiAB stabilised in amphipol A8-35 was deposited from solution (purple and ❷). hPOPC was then deposited at 20°C from a vesicle suspension where it formed a bilayer around the membrane spanning region of PqiAB by displacing solution stabilising amphipol (red and ❸). **(D)** Corresponding NR derived component volume fraction *vs.* distance profiles. The line widths in the NR model data fits depict the range of acceptable fits within the 65% confidence interval range of the posterior distributions of the fitted model parameters after MCMC error analysis. Ambiguity in the resolved surface structure is shown as line widths in the SLD profiles which are derived from the 65% confidence interval of the parameter posterior distributions. *Denotes contributions to the interfacial structure which are inferred from the fitted nSLD values.

NR confirmed that the resulting interfacial architecture is consistent with earlier assemblies (Figure 3B-D; cf. Figures 1 and 2). Importantly, maintaining the dDPPC in the gel phase suppressed inter-bilayer GPL mixing, yielding a structurally stable (“static”) system suitable for detailed analysis (Figure 3D). Quantitative modelling revealed a proximal bilayer composition of 64:36 d/h GPLs and a distal bilayer composition of ∼20:80, with an overall distal bilayer coverage of ∼40% (Table 4).

**Table 4.** A comparison of Pqi-complex separated proximal and distal dDPPC/hLipid bilayer GPL composition, thickness and coverages after deposition below the dDPPC Tm (20°C) and after heating the sample above the dDPPC Tm (42°C) so that both GPL components were in the fluid phase.

| Region | Parameter | 20 °C | 42 °C |
| --- | --- | --- | --- |
| Proximal Bilayer | bilayer thickness (Å) | 52 (49–55) | 42 (39–45) |
|  | bilayer coverage (%) | 96 (93–99) | 98 (96–100) |
|  | bilayer composition | 64 / 36 | 55 / 45 |
|  | Inner leaflet (dPPC / hLipid*) | 68 (65–71) / 32 (29–35) | 58 (54–62) / 42 (38–46) |
|  | Outer leaflet (dPPC / hLipid) | 60 (57–63) / 40 (37–43) | 52 (49–55) / 48 (44–51) |
| Inter-bilayer region | Pqi span (Å) | 250 (245–255) | 260 (255–265) |
| Distal Bilayer | bilayer thickness (Å) | 51 (44–58) | 39 (34–45) |
|  | bilayer coverage (%) | 40 (36–48) | 64 (53–77) |
|  | bilayer composition (dPPC / hLipid*) | 21 (19–23) / 79 (77–81) | 48 (46–50) / 52 (50–54) |

Similarly for the dDMPC:hPOPC proximal:distal bilayer system, we observed the presence of some protiated material (36% total material) within the proximal bilayer following system construction that we attribute to detergent and protein associated GPLs (19% total material, 52% of protiated material), the remainder was hPOPC (17% total material, 48% of protiated material), likely deposited onto exposed regions of the support directly, as only an 80% initial surface coverage was achieved in this system (see Supplementary Table S2) which increased to ∼95% on distal bilayer deposition (Table 4), thereby reducing surface defects in the proximal membrane and increasing the amount of protiated material present (see Supplementary Table S2).

The ∼20% deuterated material in the distal bilayer was likely the result of a minor population of dDPPC which transferred to the distal bilayer region during Pqi complex formation at 42℃ and after hPOPC addition.

Next, we assessed the effect of incremental temperature increases on the interfacial structure of the system. As the temperature rose above the pre-phase transition temperature of dDPPC (∼29 °C (Bryant et al. 2019)), pronounced changes were observed in the NR data, with no further significant evolution above 38 °C, where the lipid would be in its fluid phase (Figure 4A). Analysis confirmed that the overall double-bilayer architecture was retained throughout (Figure 4B-D). However, substantial GPL mixing occurred between the proximal and distal bilayers, accompanied by thinning of both bilayers due to GPL melting. This was also coupled to an increase in distal bilayer coverage, likely reflecting redistribution and equilibration of the total GPL pool following dDPPC melting and proximal bilayer thinning (see Table 4).

**Figure 4.**
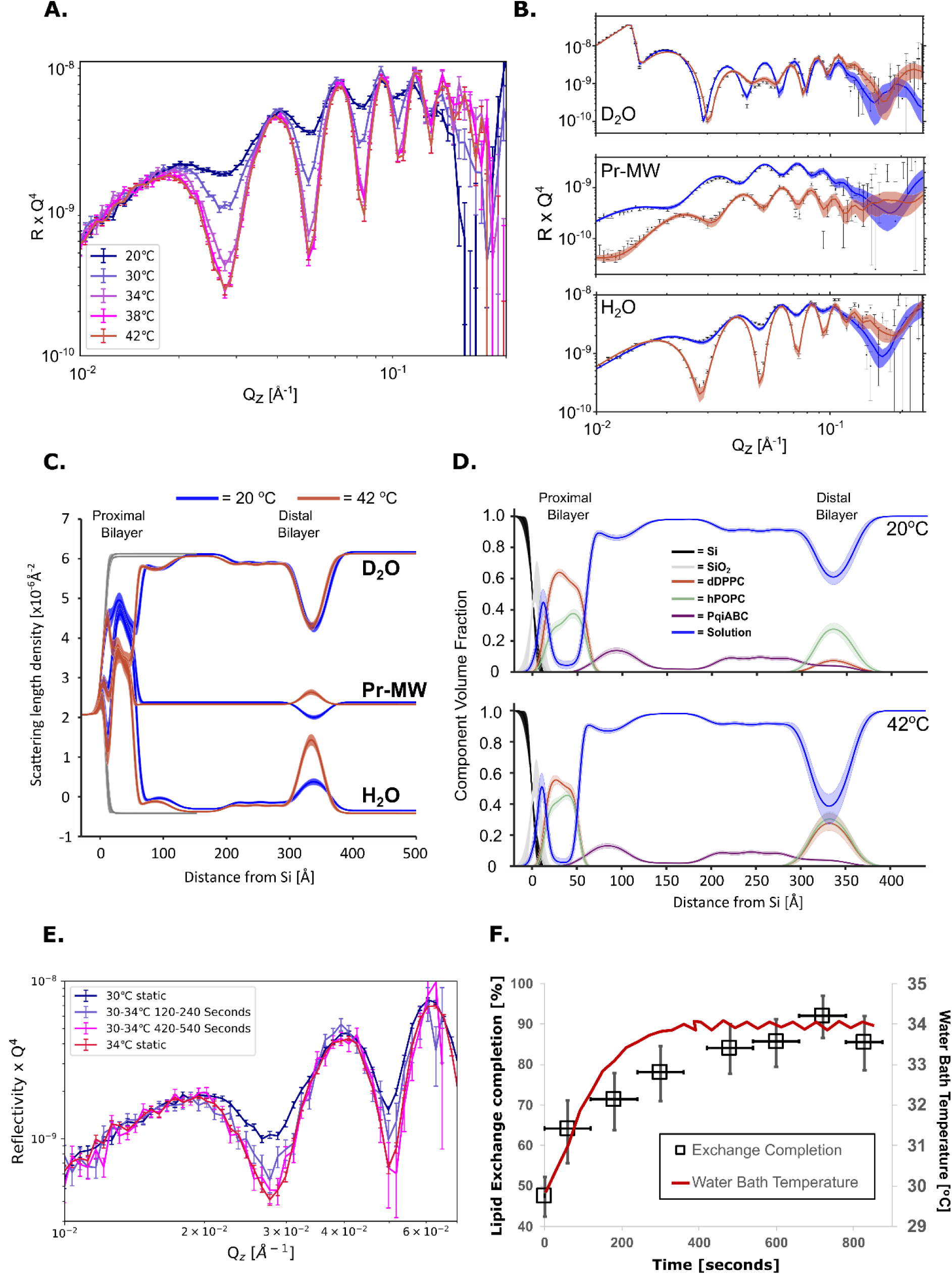
Dynamical control of Pqi-mediated intermembrane GPL transport through GPL melting. **(A)** NR profiles obtained in a H_2_O buffer solution for the Pqi complex separated double bilayer sample deposited as a proximal bilayer of dDPPC and a distal bilayer of hPOPC as the sample was incrementally increased in temperature from 20℃ to 42℃. Note how the depth of the Kiessig fringes increased from 20℃ to 38℃ due to the increased equilibration of the GPL components between the proximal and distal bilayers (this particular contrast is sensitive to the deuterated GPL content of the proximal and distal bilayers). (B) Corresponding NR profiles (error bars) and model data fits (lines) and (C) scattering length density profiles for the data observed in (A) at 20 °C, below the T_M_ of dDPPC (blue) and 42 °C (red), where both dDPPC and hPOPC were in the liquid phase. (D) The corresponding component volume fraction vs distance profiles reveal that the proximal and distal bilayers homogenised in GPL composition once the dDPPC was in the fluid phase. The temporal relationship between sample temperature and interbilayer lipid mixing was examined by comparing the depth of the 1st minima (∼0.029 Å^-1^) in the H_2_O solution contrast against time as the bilayer was heated from 30 to 34℃, i.e. where the largest change in this feature was seen (A). Time-resolved NR data (E) revealing inter-bilayer mixing was concurrent with the change in sample temperature (F), suggesting a rapid temperature-controlled process.

Prior to dDPPC melting, the outer leaflet of the proximal bilayer and the distal bilayer exhibited d/h GPL ratios of ∼65:35% and ∼20:80%, respectively. Following melting, these compositions converged to approximately ∼50:50% in both bilayers, consistent with equilibration across the system (Table 4). These observations suggest that the mixing observed previously for the dDMPC:hPOPC system (Figure 2) was not the result of direct interaction between the spatially separated bilayers, but rather mediated through PqiABC.

To further address whether the mixing could be the result of spontaneous lipid mixing across the interbilayer gap, we assessed the rate of exchange between the bilayers. Mixing of the bilayers predominantly occurred between 30-34 °C, between the pre-phase (∼29℃) and main gel-to-liquid phase transition (∼37℃) temperatures for dDPPC (Bryant et al. 2019). By time slicing the NR data during the temperature ramp, we were able to directly observe the development of bilayer mixing in real-time (Figure 4E & F). The results indicate that exchange occurred at a rate comparable to the change in the temperature of the system, with temperature induced mixing reaching completion within ∼450 seconds (Figure 4F). Previous experiments investigating spontaneous lipid transport showed 20-30% mixing of DMPS, DMPG and DMPA (shorter chain GPLs) requiring >10,000 seconds at the higher temperature of 37 °C. The rapid exchange observed, using longer chain (i.e. less transferable (Richens et al. 2017)) lipids and at lower temperatures, therefore casts serious doubt on the observed exchange being spontaneous. Rather the speed of exchange further supports PqiABC mediated transport.

Finally, to confirm that the changes in movement observed were the result of phase transition, we repeated the dDPPC:hPOPC system deposition (see Supplementary, Section 2) which revealed the same trends in proximal-to-distal bilayer lipid equilibration upon melting of the proximal bilayer dDPPC.

### Fluorescence measurements support a role for PqiABC in glycerophospholipid transport

To complement the NR studies and provide an orthogonal approach to probe GPL transport, we developed an *in vitro* FRET-based assay. In this assay, purified DDM-solubilised PqiABC, either expressed as the intact complex or reconstituted from its subcomplexes (PqiAB and PqiC), was diluted into a 1:1 mixture of unlabelled POPC liposomes and liposomes containing the fluorescent GPLs nitrobenzoxadiazole-PE (NBD-PE) and rhodamine-PE (Rhod-PE). The final DDM concentration was maintained below the level required to destabilise or solubilise liposomes (Figure 5A), preserving vesicle integrity while favouring partitioning of PqiABC into the lipid bilayer, enabling formation of PqiABC conduits between liposomes rather than remaining within the detergent micelle. GPL transport was monitored via changes in NBD fluorescence: initially quenched by proximity of Rhod-PE via FRET, NBD fluorescence increased upon GPL transport as dilution of the fluorophores reduced quenching, resulting in an increased emission at 535 nm (Figure 5A).

**Figure 5.**
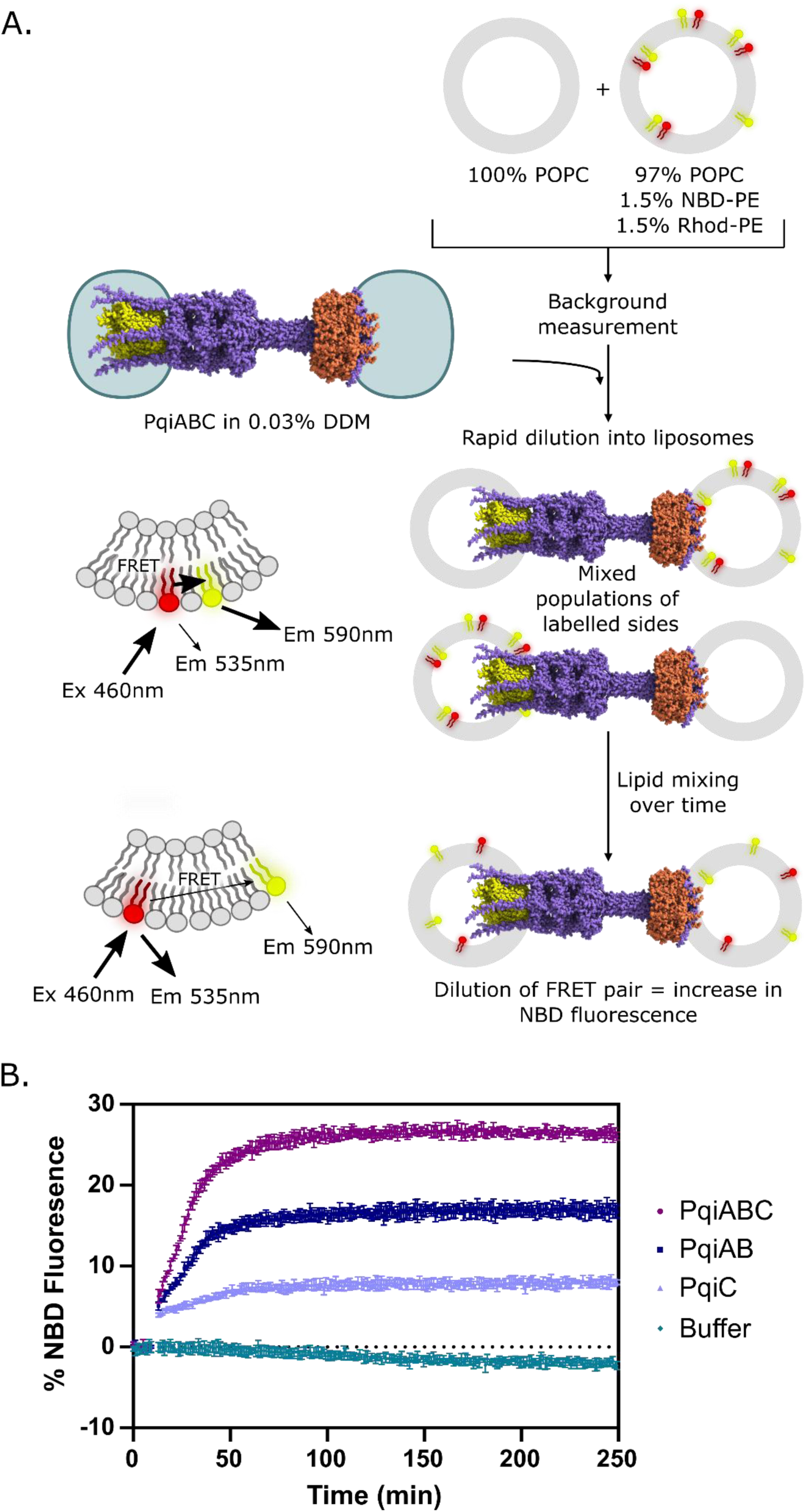
FRET-based assay to monitor glycerophospholipid (GPL) transport by PqiABC. **(A)** Schematic of the FRET assay. POPC liposomes and POPC liposomes containing NBD-PE and Rhod-PE (POPC:NBD-PE:Rhod-PE) were mixed in equimolar amounts with buffer in a 96-well plate, and fluorescence was monitored until a stable baseline was established. DDM-solubilised protein was then added such that the final DDM concentration was below the level required to destabilise or solubilise liposomes. NBD fluorescence was measured continuously (excitation, 460 nm; emission, 535 nm). Initially, NBD fluorescence is quenched through Förster resonance energy transfer (FRET) to the nearby Rhod-PE acceptor. As GPLs redistribute between liposomes, the fluorophores become diluted, reducing FRET efficiency and resulting in dequenching of NBD fluorescence, observed as an increase in emission at 535 nm. Arrow thickness represents the relative efficiency of energy transfer. **(B)** Representative FRET assay showing the increase in NBD fluorescence over time. Fluorescence values were normalised to the initial baseline and maximal fluorescence. Protein was added at a final concentration of 1 µM. Data are presented as mean ± s.d. (n = 3 independent replicates).

Figure 5B shows the time-dependent change in NBD fluorescence for PqiABC. Signals were normalised to a liposome only control and scaled using detergent-mediated solubilisation to define the maximal fluorescence (corresponding to complete fluorophore redistribution). A pronounced increase in fluorescence was observed over time in the presence of PqiABC, consistent with GPL transport between membranes and in agreement with the NR data.

To assess the contribution of individual components, subcomplexes were tested separately. Removal of parts of the Pqi complex reduced the overall fluorescence increase (Figure 4B), indicating diminished GPL transport. PqiC alone showed minimal activity (∼7% increase over 250 min), whereas PqiAB displayed modest activity (∼16%), in contrast to the substantially higher activity observed for PqiABC (∼25%). Overall, these observations align with complementation assays demonstrating that all components of the PqiABC system are required for full function in vivo (Supplementary Figure S6). The differing activities of the subcomplexes indicate that PqiAB plays a more critical role in GPL transport than PqiC. This is consistent with our previous structural model in which PqiC acts primarily as a stabilising outer membrane anchor for the PqiB needle (Cooper et al. 2024). Loss of this anchor would be expected to reduce the stability of needle–membrane interactions while still permitting transient contacts capable of supporting low levels of transport. By contrast, the absence of PqiAB essentially abolishes transport by removing the conduit itself. The residual signal observed in the PqiC-only sample is therefore likely to reflect background lipid exchange or membrane interactions arising from PqiC binding to the liposome surface, rather than genuine protein-mediated translocation. Finally, the distinct transport profiles of the different protein assemblies provide strong evidence that the fluorescence changes arise from protein-mediated GPL transport rather than spontaneous lipid exchange, which would be expected to occur at similar rates regardless of the protein composition.

### Conclusions

Collectively, our structural and functional analyses establish PqiABC as an envelope-spanning GPL transport system that directly mediates GPL exchange between the inner and outer membranes. NR reveals that PqiABC forms a continuous conduit capable of supporting bidirectional GPL equilibration across a physiologically relevant periplasmic distance, while phase-controlled experiments demonstrate that this process is dependent on membrane fluidity and does not arise from membrane contact or fusion. These findings are further supported by fluorescence-based assays, which confirm that GPL transport occurs through the intact PqiABC complex and not a result of spontaneous lipid exchange. While our *in vitro* data are most consistent with a passive equilibration mechanism, we cannot exclude the possibility that, *in vivo*, PqiABC-mediated transport may be coupled to cellular energy sources, such as the proton motive force, to impart directionality under specific physiological conditions.

When considered alongside the well-established upregulation of the *pqi* operon in response to envelope stress, our findings support a model in which PqiABC functions as a dynamic GPL redistribution pathway that helps preserve envelope integrity under damaging conditions. We propose that PqiABC facilitates the passive equilibration of GPLs between the inner and outer membranes, enabling redistribution of the membrane lipid pool to maintain GPL homeostasis. Such bidirectional transport could promote the dilution or replacement of locally damaged GPLs while preventing imbalances in membrane composition that compromise barrier function. In this context, PqiABC may act as a stress-responsive GPL equilibration conduit that buffers the Gram-negative envelope against oxidative and other envelope stresses, thereby enhancing membrane resilience and promoting cell survival.

As part of our QCM-D and NR studies into the PqiABC lipid transport system, we developed an *in situ* self-assembled model of the complete Gram-negative envelope. The resulting interfacial architecture comprises inner and outer membranes separated by a spatially accurate periplasmic region. To our knowledge, no previous model has reproduced the compositional and structural complexity of the entire diderm envelope in a single experimentally accessible system.

The three-step assembly protocol enables the generation of this architecture directly within the measurement environment, facilitating its application across a variety of analytical techniques. As a result, the platform provides a versatile assay for the investigation of intermembrane transport processes and establishes a new experimental framework for molecular-level studies of Gram-negative envelope biochemistry.

## CRediT authorship contribution statement

**Hannah E. Johnston:** Conceptualisation, Methodology, Investigation, Writing - original draft.

**Luke A. Clifton:** Conceptualisation, Methodology, Software, Formal analysis, Investigation, Writing - original draft, Writing - review & editing, Visualisation.

**Charlotte B. Wilson:** Conceptualisation, Methodology, Validation, Investigation, Writing - original draft

**Pooja Sridhar:** Methodology, Validation, Investigation

**Abdulaziz Alzahrani:** Validation, Investigation

**Adam Colyer:** Investigation

**Richard Logan:** Investigation

**Stephen C.L. Hall**: Resources

**David J. Hardy:** Investigation

**Timothy J. Knowles:** Conceptualisation, Methodology, Investigation, Resources, Writing - original draft, Writing - review & editing, Visualisation, Supervision, Project administration, Funding acquisition.

## Acknowledgements

LAC would like to thank Dr Max Skoda (ISIS, RAL) and Dr Alessandra Luchini (University of Perugia) for helpful discussions.

## Supporting Information and Data Availability

Section 1 of the supporting information provides a table of oligonucleotide primers used in this study, figures detailing the process used to convert PqiC and PqiABC structures into the 1-D density profiles used in NR analysis, tables of the nSLD values used in NR data fitting and the analysis of a dDMPC:PqiC SLB with reduced DDM contamination. Section 2 describes repeat NR measurements on temperature-controlled bilayer phospholipid mixing studies on a Pqi-spanning double bilayer sample with a proximal deposited as dDPPC and a distal bilayer deposited as hPOPC. Section 3 of the supporting information *In vivo* assays on Pqi function. Section 4 provides fitted and fitting derived parameter tables from NR data analysis. NR data, custom rascal models used to analyze the data, fluorescence and QCM-D data are available on Zenodo: DOI 10.5281/zenodo.21507815

