## Supplementary Information for "PqiABC forms a membrane-bridging conduit and mediates bidirectional phospholipid transport across the Gram-negative bacterial envelope"

##### **Contents:**

**Section 1:** Supplementary Figures and Results to support the neutron reflectometry measurements and analysis in the main article

Table S1. Primers used in this study

Table S2. Table of component Scattering Length density values used in NR data fitting.

Figures S1 and S2. Generation of the 1-dimensional density profiles of PqiC and PqiABC used in NR data fitting.

Table S3. Structural parameters obtained for the initial stages (1 and 2) of the assembly of a PqiABC spanning double bilayer sample with a proximal bilayer deposited as dPPC and a distal bilayer deposited as hPOPC.

Figure S3. NR data, SLD profiles and component volume fraction profiles for a dDMPC bilayer with a outer leaflet surface population of PqiC deposited from 1:5 (w/w) protein-lipid vesicles showing a lower DDM contaminant than the protein-lipid SLBs deposited from 1:2.5 (w/w) vesicles shown in the main manuscript.

**Section 2:** Additional results showing repeat measurements on the assembly of a Pqi complex spanning double bilayer sample system with a proximal bilayer deposited as dPPC and a distal bilayer of hPOPC and measurements on the temperature-controlled mixing between the proximal and distal bilayers.

Figure S4 and S5. Additional results for NR measurements on the assembly (Fig S4) and temperature controlled proximal and distal bilayer mixing (Fig S5).

**Section 3:** Additional Results : *In vivo* assays.

Figure S6. Phenotypic complementation assay assessing the contribution of Pqi components to survival.

**Section 4:** Tables (S5-18) of fitted and fitting derived parameters from model-to-data fits of NR data sets shown in the main manuscript and supporting information section 2.

### Section 1: Supporting Figures and Tables.

Supplementary Table 1. Oligonucleotide primers used in this study

|  | Primer |  |
| --- | --- | --- |
| Construct | Forward | Reverse |
| PqiC-ΔC-His | TGAGATCCGGCTGCTAAC | AGGTAGACGCTTTATCTCTTG |
| PqiA N-strep | CAGTTCGAAAAATGCGAACAT<br>CATCATGCC<br>CGGGTGGCTCCACATATGTAT<br>AT | CTCCTTCTTAAAGTTAAAC |

**Supplementary Table 2. Neutron Scattering Length Density Values used in NR Fitting.**

| Component | nSLD Value [$\times 10^{-6} \text{ \AA}^{-2}$] | Source |
| --- | --- | --- |
| Phosphatidylcholine (PC) head-group | 1.98 | (John et al. 2021) |
| d-DMPC Tails Gel | 7.23 | (Browning et al. 2017) |
| d-DMPC Tails Liquid | 6.55 | (Browning et al. 2017) assuming 5% H contamination on chains |
| d-DPPC Tails Gel | 7.43 | (Carrascosa-Tejedor et al. 2020) |
| d-DPPC Tails Liquid | 6.72 | Derived from (Carrascosa-Tejedor et al. 2020) assuming tail vol is $0.825 \text{ nm}^3$ in gel phase and $0.913 \text{ nm}^3$ in fluid phase; |
| h-POPC Tails | -0.30 | (John et al. 2021) |
| Amphipol A8-35 | 1.06 | (Arunmanee et al. 2014) |
| PqiC in D<sub>2</sub>O buffer | 1.90 | Calculated:<br>http://psldc.isis.rl.ac.uk/Psldc/ |
| PqiC in H<sub>2</sub>O buffer | 3.24 | Calculated:<br>http://psldc.isis.rl.ac.uk/Psldc/ |
| PqiABC in D<sub>2</sub>O buffer | 3.16 | Calculated:<br>http://psldc.isis.rl.ac.uk/Psldc/ |
| PqiABC in H<sub>2</sub>O buffer | 1.85 | Calculated:<br>http://psldc.isis.rl.ac.uk/Psldc/ |
| DDM Heads | 1.31 | (Vacklin et al. 2005) |
| DDM Tails | -0.39 | (Vacklin et al. 2005) |
| *E. coli* phospholipid tails | -0.55 | (Lind et al. 2015) |
| *E. coli* phospholipid heads in H<sub>2</sub>O | 1.55 | (Lind et al. 2015) |
| *E. coli* phospholipid heads in D<sub>2</sub>O | 2.16 | (Lind et al. 2015) |
| *E. coli* phospholipid average | 0.1 | Calculated from values in (Lind et al. 2015) |

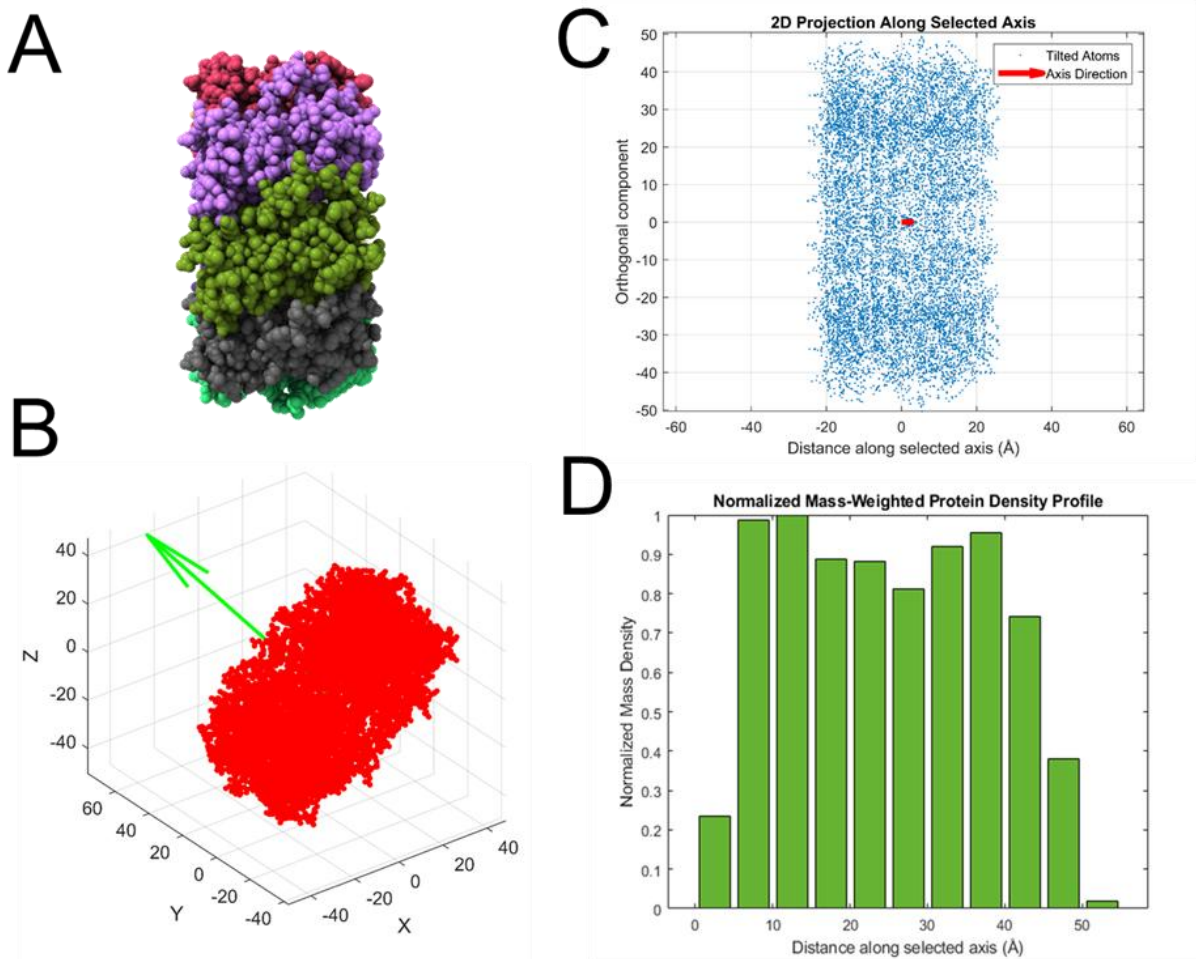

**Supplementary Figure S1 - (A)** The proposed structure of the PqiC octamer used in NR analysis. **(B)** The projection of the PqiC octamer in three-dimensions, with the minor axis used to determine the 1-D density profile of this protein complex shown via a green arrow. **(C)** The resulting one-dimensional density profile of the PqiC octamer along its minor axis. **(D)** Plot of this structure in two dimensions to compare with the features of the density profile.

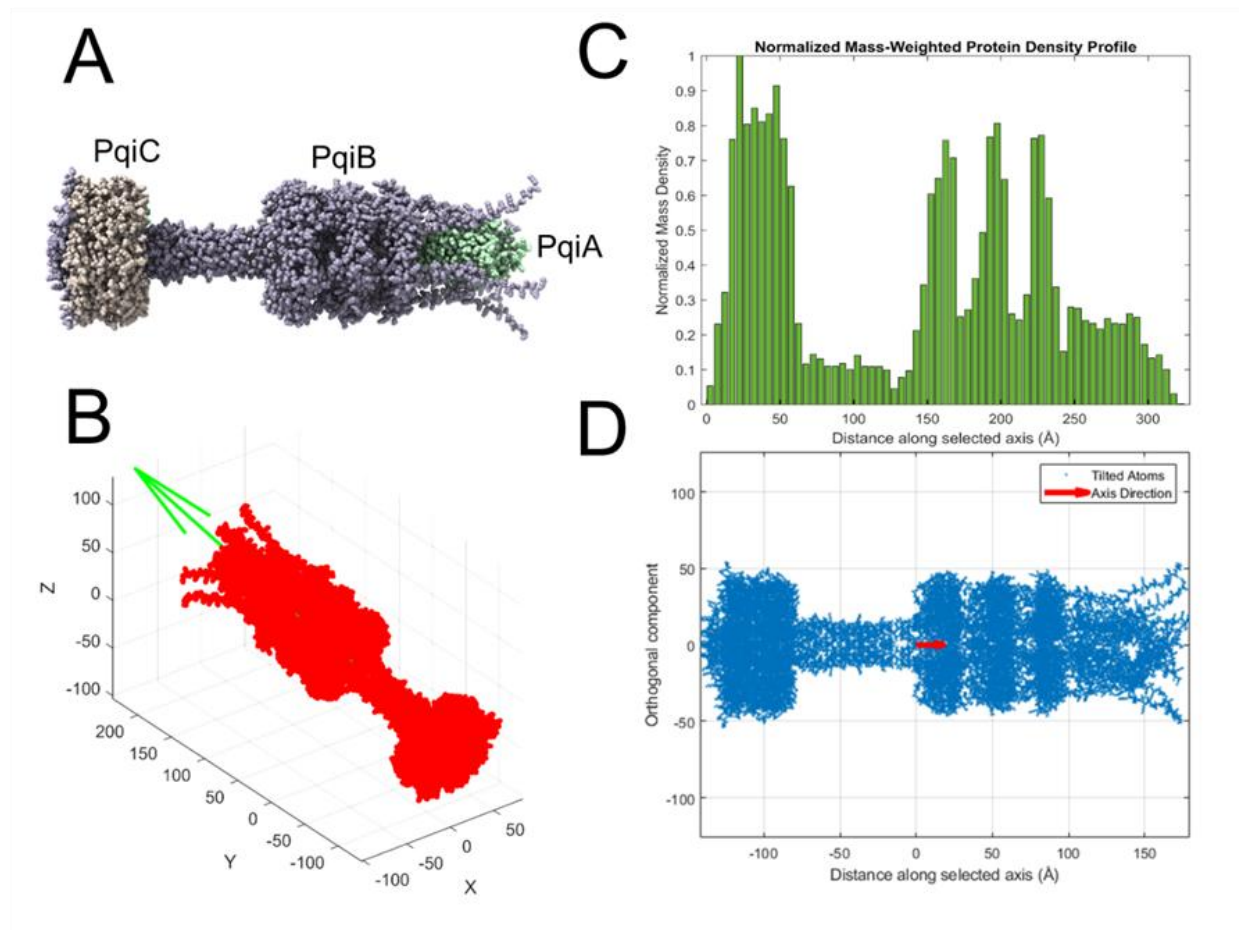

**Supplementary Figure S2 - (A)** The proposed structure of the PqiABC complex used in NR analysis. The projection of this structure in three-dimensions **(B, red)** with the major axis used to determine the 1-D density profile of this protein complex shown **(B, green)**. The resulting one-dimensional density profile of the Pqi complex along its major axis **(C)** below a 2D projection of all the atoms in the x and y directions are plotted against the z-direction **(D)** to compare with the features of the density profile derived from this.

**Supplementary Table 3: NR-fitting derived structural parameters obtained for the initial stages (1 and 2) of the assembly of a PqiABC spanning double bilayer sample with a proximal bilayer deposited as dPPC and a distal bilayer deposited as hPOPC. NR Data, fits, nSLD and volume fraction profiles shown in Figure 3 of the main manuscript.**

| **Fabrication Stage** | **Proximal SLB Thickness** | **Proximal SLB Composition** | **Pqi Coverage** | **SLB to Amphipol Distance** | **Amphipol region Coverage** |
| --- | --- | --- | --- | --- | --- |
| **1.**<br>**dPPC-PqiC SLB** | 47 Å<br>(43, 50) | **dPPC:**<br>69% (65, 74)<br>**Contaminant (likely DDM):**<br>13% (9, 17)<br>**Solution:**<br>18% (9, 26) | **PqiC:**<br>14%<br>(11, 17) | N/A | N/A |
| **2.**<br>**dPPC-PqiABC -Amphipol** | 49 Å<br>(45, 52) | **dPPC (average):**<br>68% (61, 75)<br>**Contaminant (likely DDM and native lipids):**<br>14% (8, 25)<br>**Solution:**<br>18% (0, 31) | **PqiABC:**<br>17%<br>(15, 19) | 260Å<br>(250, 270) | **Native Lipid:**<br>7% (5, 11)<br>**dPPC:**<br>5% (3, 8) |

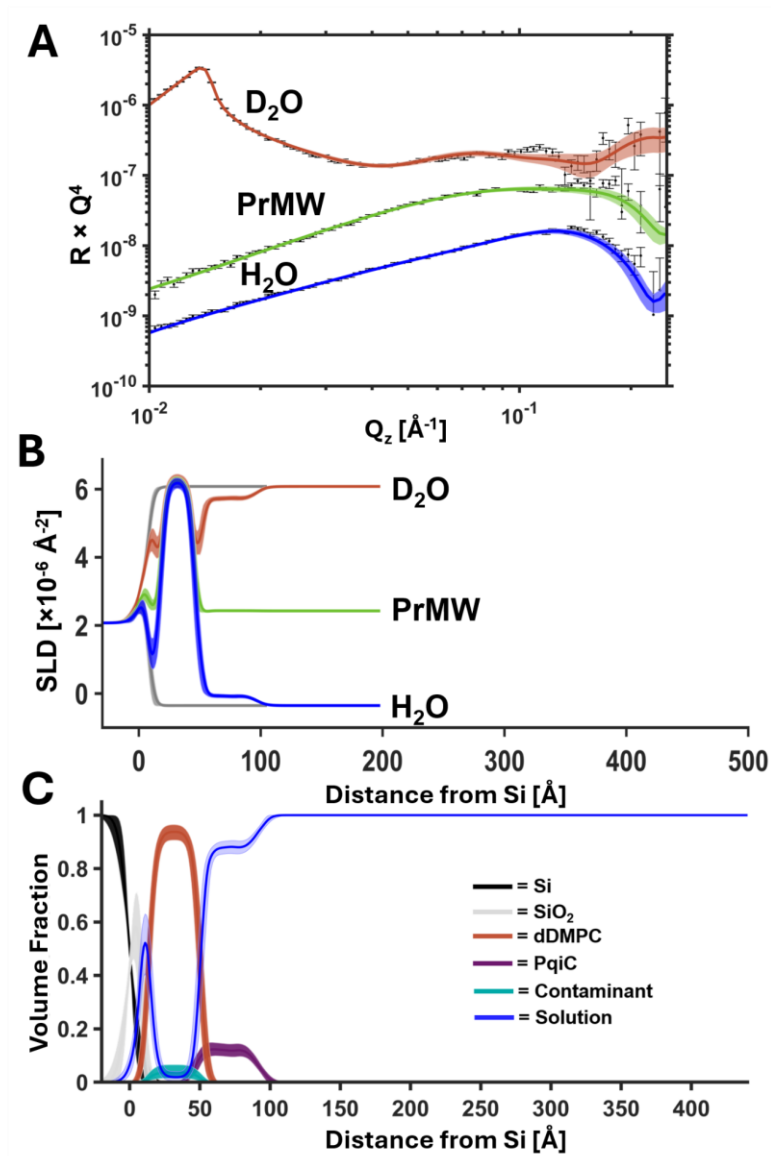

**Supplementary Figure S3 - NR data (A, error bars) and model data fits (A, lines) and the scattering length density profiles (B) and component volume fraction vs distance profiles these fits represent (C) for a dDMPC supported lipid bilayer with PqiC bound on its outer leaflet surface deposited from 1:5 (w/w) protein-lipid vesicles. Note that significantly less contaminant (C, identified as DDM surfactant) is found in this sample compared to samples deposited from 1:2.5 (w/w) protein-lipid vesicles.**

### Section 2: Additional Results and Discussion

#### Repeat NR measurements monitoring Pqi mediated lipid transport using Gel-to-liquid phase as an activity switch.

The assembly of a PqiABC mediated double membrane spanning system with a proximal dPPC bilayer and a distal hPOPC bilayer was structurally examined using NR. This was a repeat measurement to that described in the main manuscript (Figures 3 and 4). Temperature was used to control the phase of the lipid components during the distal (h-)bilayer deposition and thus control inter-bilayer lipid mixing. This was conducted with an aim to produce a bilayer where proximal and distal bilayer mixing was initially prevented via keeping the dPPC in the gel phase while the hPOPC was deposited onto the distal bilayer region of the structure.

Figure S4 shows the neutron reflectometry data, model-to-data fits and SLD profiles obtained during the assembly of this sample. The assembly process showed both a relatively high coverage of PqiC on the surface of the dPPC SLB (~14%) resulting in a high coverage of PqiABC after the assembly of the full protein complex of ~18%.

During the assembly of the full Pqi complex, via the addition of amphipol-stabilized Pqi-AB to a dPPC SLB with an outer leaflet population of PqiC, the temperature of the dPPC was kept in the gel phase (at 20°C). In all other experiments (as described in the main manuscript) the proximal bilayer lipids were in the fluid phase during this stage of the Pqi complex-mediated double bilayer assembly. Interestingly, analysis of the resulting NR data revealed that only an ambiguous, low coverage (0-2%) of amphipol could be resolved around the hydrophobic belt of PqiAB (see Figs S4 B and C). Indeed, when this data was compared to samples where the proximal bilayer lipids had been in the fluid phase during full Pqi complex assembly the distinct high frequency fringes visible in the NR data (See Figures 1-3) were not apparent (see Figure S4). From this comparison we concluded that the material observed around the hydrophobic belt of PqiAB observed during the Pqi-assembly stage for the samples described in the manuscript (noted as stage II in Figs 1-3) were a mixture of native *E. coli* lipids carried internally by Pqi from purification onwards and proximal bilayer lipids which formed low coverage lipid bilayers due to passive transport through PqiABC even in the absence of the distal bilayer.

The sample then held at 28°C during hPOPC addition to form a high coverage distal bilayer. Some mixing of hPOPC was observed into the predominantly dPPC proximal bilayer (Figure S4, B and C, III) likely through the deposition of hPOPC vesicles into the ~20% coverage of proximal bilayer defects.

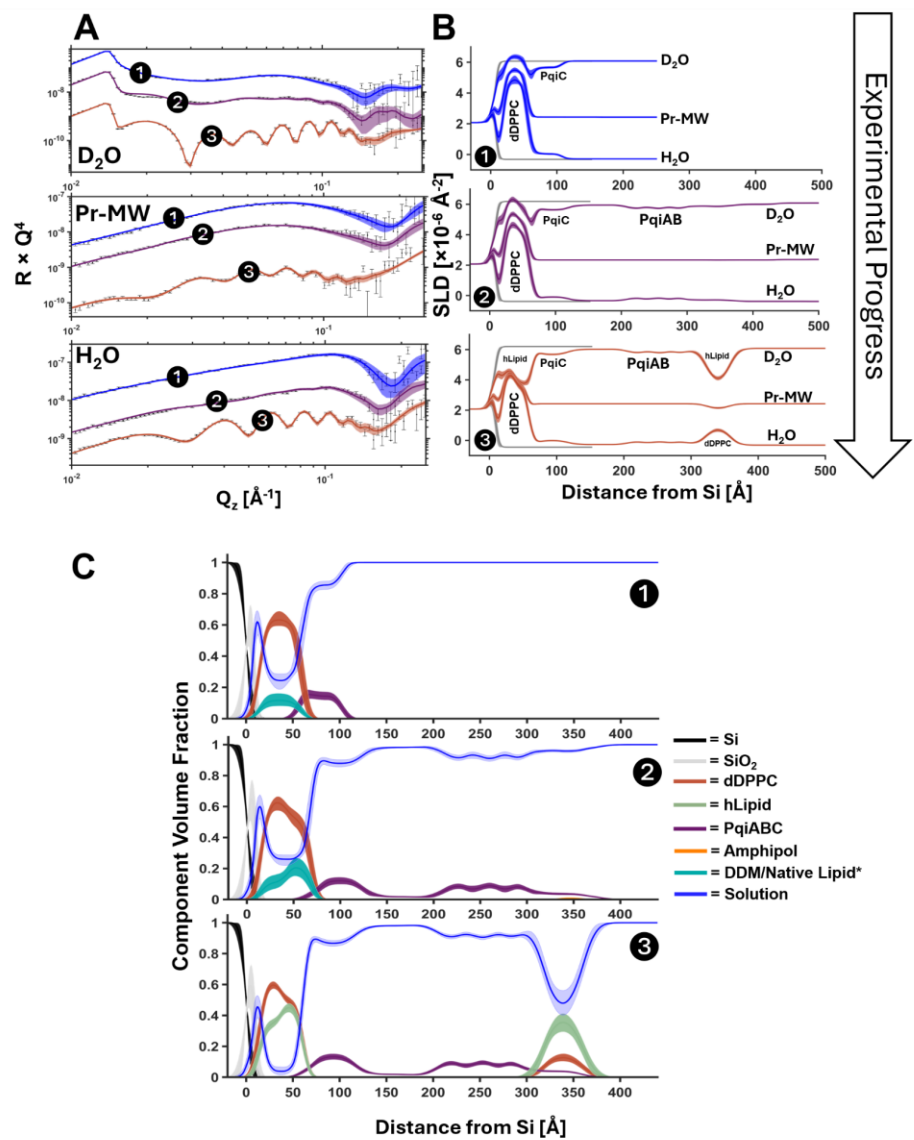

**Supplementary Figure S4** - Neutron reflectometry profiles (**error bars, A**), model data fits (**lines, A**), scattering length profiles (**B**) and component volume fraction profiles these fits describe (**C**) for the step-wise assembly of a Pqi-complex spanning double bilayer sample with dDPPC deposited as the proximal bilayer at 42°C (**1**) and hPOPC deposited as the distal bilayer at 28°C (**3**). The assembly of the full Pqi complex (C **2**) was undertaken at 20°C with the dDPPC proximal bilayer in the gel phase. It was noted that the lipid distributions seen after Pqi complex assembly when the proximal bilayer was in the fluid phase were not present, suggesting these were lipid as opposed to amphipol. Amphipol was only ambiguously resolved around the hydrophobic region of PqiAB in this data (**2**). \*Denotes contributions to the interfacial structure which are inferred from the fitted nSLD values.

After the addition of hPOPC to the sample at 28°C the sample was heated to 42°C and the change in the distribution of components across the Pqi-spanning double bilayer structure was examined. NR analysis (Figure S5) revealed that as the sample was heated from 28°C to 42°C in 3°C steps mixing between the proximal and distal bilayers occurred (Figure S5B). Notably, after dPPC melting there was an increase in dPPC in the distal bilayer (Figure S5B and C and Table S3) and hydrogenous lipid in the proximal bilayer, likely due to Pqi facilitated transport of the fluid phase lipid across the ~260 Å gap between the proximal and distal bilayers (see Table S3).

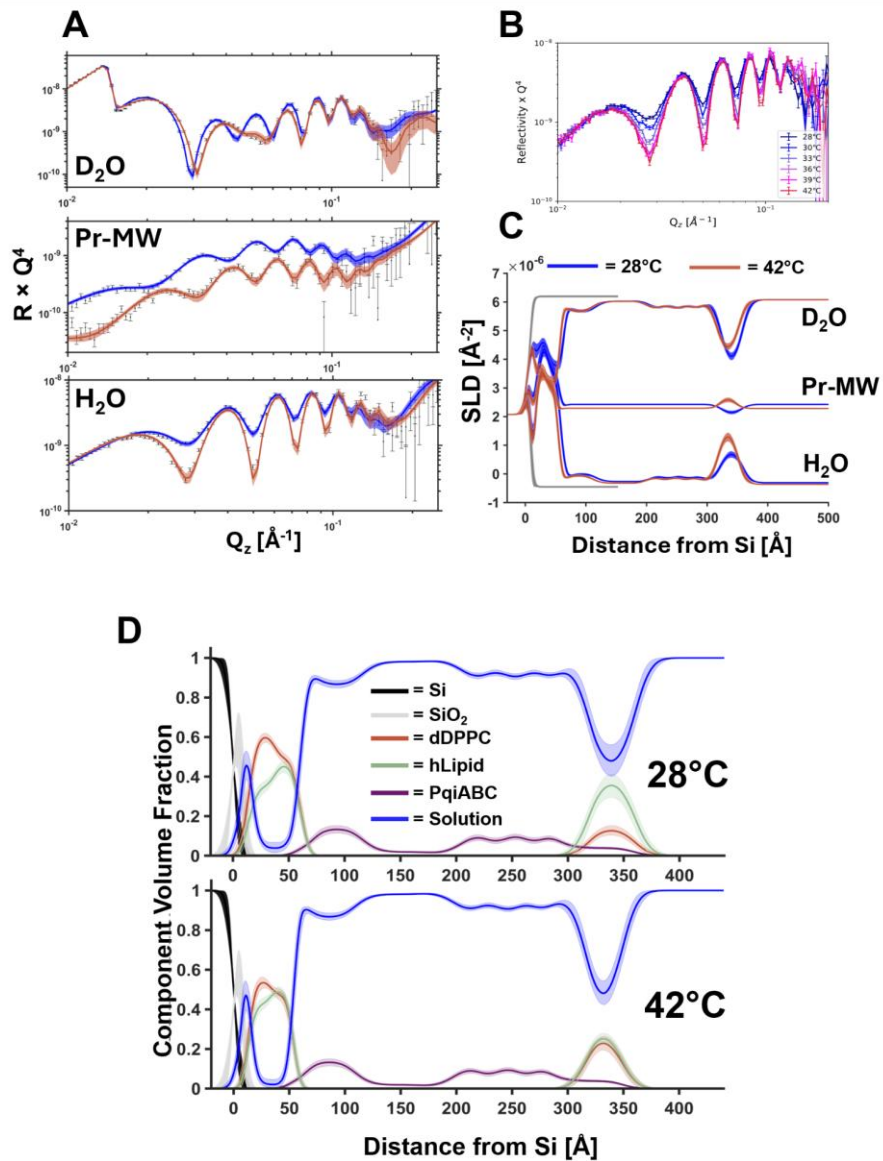

**Supplementary Figure S5 - Repeat dynamical control of Pqi mediated intermembrane lipid transport through phospholipid melting.** Neutron reflectometry profiles (**A**, error bars) and model data fits (**A**, lines) and the scattering length density profiles these fits describe (**B**) for a double lipid bilayer on a Pqi scaffold with a proximal bilayer deposited as dPPC and a distal bilayer deposited partially as hPOPC after deposition at 28°C, below the T<sub>m</sub> of dPPC (blue) and after heating to 42°C (red), where both the dPPC and hPOPC were in the liquid phase. The change in the experimental NR data from the sample in the H<sub>2</sub>O buffer contrast during every 3°C increments between 28°C and 42°C is shown (**B**). The fitting derived component volume fraction vs. distance profiles for the samples at the two temperatures are shown (**C**) revealing the equilibration of the proximal and distal bilayer components above the T<sub>m</sub> of dPPC.

The coverage of lipid on the distal bilayer increased by ~6% after dPPC melting while the thickness of both the proximal and distal bilayers was reduced (see Table S4). This h/d lipid composition of both bilayers reached ~equilibrium after melting, which was especially noted when comparing the outer leaflet of the proximal bilayer with the distal bilayer lipid composition. This data therefore further suggests that PqiABC provides a means of bi-directional lipid transfer between the spatially separated proximal and distal bilayers. This in turn provides further evidence that the Pqi complex serves as a conduit that enables equilibration of lipid contents between the inner and outer membranes of Gram-negative bacteria.

**Supplementary Table 4. A comparison of Pqi-complex separated proximal and distal dPPC/hLipid bilayer GPL composition, thickness and coverages after deposition below the dPPC T<sub>m</sub> (28°C) and after heating the sample above the dPPC T<sub>m</sub> (42°C) so that both GPL components were in the fluid phase. \*hLipid is a combination of hPOPC, *E.coli* native lipids and a minor DDM contamination.**

| Region | Parameter | 28 °C | 42 °C |
| --- | --- | --- | --- |
| Proximal Bilayer | bilayer thickness (Å) | 50 (47–53) | 44 (41–45) |
|  | bilayer coverage (%) | 96 (93–98) | 98 (95–100) |
|  | bilayer composition | 58 / 42 | 52 / 48 |
|  | Inner leaflet (dPPC / hLipid*) | 64 (61–67) / 42 (41–43) | 56 (53–59) / 44 (41–47) |
|  | Outer leaflet (dPPC / hLipid*) | 51 (49–54) / 49 (46–51) | 49 (46–52) / 51 (48–54) |
| Inter-bilayer region | Pqi span (Å) | 260 (255–265) | 260 (255–265) |
| Distal Bilayer | bilayer thickness (Å) | 39 (34–46) | 35 (32–40) |
|  | bilayer coverage (%) | 53 (42–62) | 59 (49–67) |
|  | bilayer composition (dPPC / hLipid*) | 27 (25–29) / 73 (71–75) | 47 (45–49) / 53 (51–55) |

#### Section 3: Additional Results - *In vivo* assays

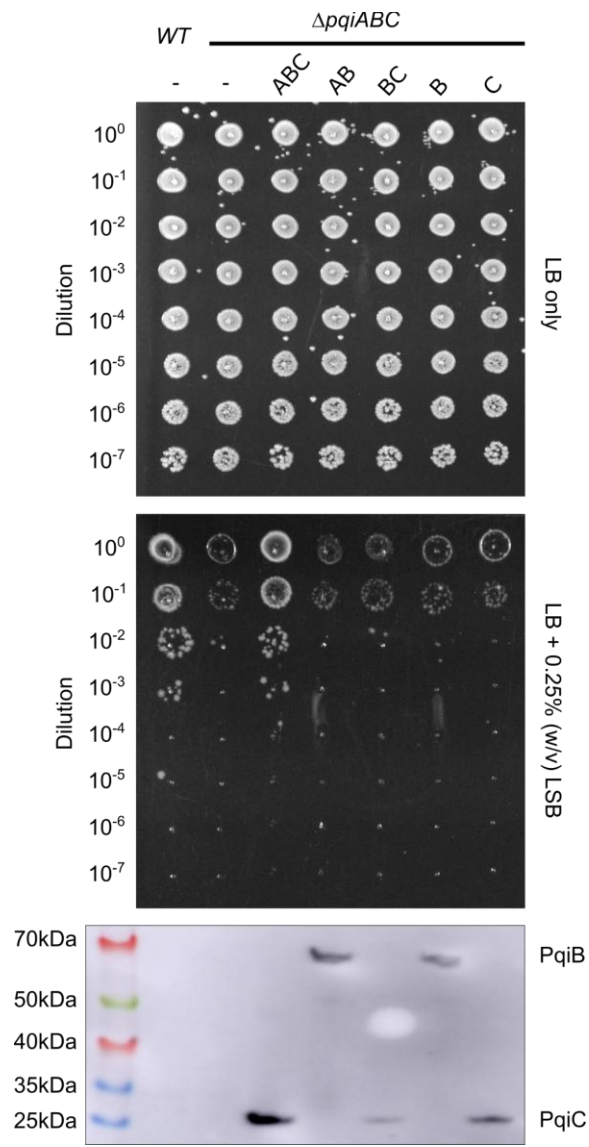

**Supplementary Figure S6 – Phenotypic complementation assay assessing the contribution of Pqi components to survival in the presence of lauryl sulfobetaine.** Growth phenotypes of the indicated knockout and complemented strains are shown under detergent stress conditions. The lower panel presents a corresponding Western blot confirming stable expression of Pqi components across strains. Detection was performed using an anti-His primary antibody targeting His-tagged PqiB or PqiC.

### Section 4: NR Fitting Tables and Derived Structural Parameters

**Supplementary Table 5. Fitted and fitting derived structural parameters from POPC supported lipid bilayer at the silicon-water interface with population of PqiC bound on its outer (solution facing) surface (data and fits shown in main article Figure 1).**

| Parameter | Mean | Relative_Low_Error | Relative_High_Error |
| --- | --- | --- | --- |
| **Fitted Parameters** |  |  |  |
| {'Substrate_Roughness / Å'} | 5.5079 | 2.2613 | 2.7022 |
| {'SiO2_thickness / Å'} | 8.4654 | 1.9167 | 2.1856 |
| {'SiO2_roughness / Å'} | 5.6716 | 1.7518 | 1.3298 |
| {'SiO2_hydration / %' } | 18.696 | 10.531 | 7.6854 |
| {'Lipid_APM / Å<sup>2</sup>'} | 58.84 | 2.0992 | 2.2367 |
| {'Prox Bilayer_Coverage'} | 0.94072 | 0.032985 | 0.035786 |
| {'Prox Bilayer_HG_Hydration'} | 0.090827 | 0.062877 | 0.10958 |
| {'Prox Bilayer_Roughness / Å'} | 5.0696 | 1.2592 | 1.2017 |
| {'Water_thickness / Å'} | 5.0073 | 1.5698 | 1.3983 |
| {'PqiC Coverage'} | 0.04761 | 0.028701 | 0.033875 |
| {'PqiC roughness / Å' } | 4.671 | 3.066 | 2.9656 |
| **Derived Structural Parameters** |  |  |  |
| {'Prox Bilayer Coverage / %'} | 94.072 | 3.2985 | 3.5786 |
| {'Prox Bilayer HG thickness / Å'} | 5.4537 | 0.19974 | 0.20178 |
| {'Prox Bilayer Inner Tails thickness / Å'} | 14.984 | 0.54878 | 0.55436 |
| {'Prox Bilayer Outer Tails thickness / Å'} | 14.984 | 0.54878 | 0.55436 |
| {'PqiC Total Thickness / Å' } | 50 | 0 | 0 |

**Supplementary Table 6. Fitted and fitting derived structural parameters from proximal POPC supported lipid bilayer at the silicon-water interface with surface bound PqiABC on its outer (solution facing) surface with a low coverage distal bilayer which formed during Pqi complex assembly (data and fits shown in main article Figure 1).**

| Parameter | Mean | Relative_Low_Error | Relative_High_Error |
| --- | --- | --- | --- |
| **Fitted Parameters** |  |  |  |
| {'Substrate_Roughness / Å' } | 4.6722 | 1.7869 | 2.7993 |
| {'SiO2_thickness / Å'} | 11.58 | 2.0395 | 2.1185 |
| {'SiO2_roughness / Å'} | 4.6511 | 1.6138 | 1.4109 |
| {'SiO2_hydration / %'} | 21.506 | 10.435 | 8.886 |
| {'Lipid_APM / Å<sup>2</sup>'} | 58.218 | 1.8587 | 1.8849 |
| {'Proximal Bilayer_Coverage'} | 0.97652 | 0.019488 | 0.015358 |
| {'Proximal Bilayer_HG_Hydration'} | 0.15064 | 0.095954 | 0.12872 |
| {'Proximal Bilayer_Roughness / Å' } | 3.8605 | 0.56029 | 0.78515 |
| {'Water_thickness / Å'} | 3.7238 | 1.8177 | 1.771 |
| {'PqiABC Coverage'} | 0.095809 | 0.024363 | 0.02295 |
| {'Distal thickness / Å'} | 17.209 | 7.3743 | 11.039 |
| {'Distal coverage'} | 0.12816 | 0.050102 | 0.089101 |
| {'Distal position'} | 0.82529 | 0.024388 | 0.022936 |
| {'PqiABC roughness'} | 10.285 | 5.238 | 3.1816 |
| **Derived Structural Parameters** |  |  |  |
| {'Proximal Bilayer Coverage / %' } | 97.652 | 1.9488 | 1.5358 |
| {'Proximal Bilayer HG thickness / Å' } | 5.512 | 0.17288 | 0.18178 |
| {'ProxBilayer Inner Tails thickness / Å'} | 15.144 | 0.47497 | 0.49941 |
| {'Proxi Bilayer Outer Tails thickness / Å'} | 15.144 | 0.47497 | 0.49941 |
| {'PqiABC Total Thickness / Å'} | 320 | 0 | 0 |
| {'Prox_Bilayer_to_Distal Distance / Å'} | 265.73 | 7.8462 | 7.4161 |

**Supplementary Table 7. Fitted and fitting derived structural parameters from double POPC bilayer sample on a PqiABC scaffold (data and fits shown in main article Figure 1).**

| Parameter | Mean | Relative_Low_Error | Relative_High_Error |
| --- | --- | --- | --- |
| **Fitted Parameters** |  |  |  |
| {'Substrate_Roughness / Å'} | 4.3699 | 1.5945 | 2.2402 |
| {'SiO2_thickness / Å'} | 11.416 | 1.7794 | 1.664 |
| {'SiO2_roughness / Å'} | 3.9869 | 1.2739 | 1.6029 |
| {'SiO2_hydration / %'} | 21.093 | 9.4274 | 8.1016 |
| {'Water_thickness / Å'} | 4.3247 | 1.6298 | 1.1104 |
| {'Proximal Lipid_APM / Å<sup>2</sup>'} | 57.771 | 2.0625 | 1.962 |
| {'Proximal Bilayer_Coverage'} | 0.97377 | 0.020892 | 0.01605 |
| {'Proximal Bilayer_HG_Hydration'} | 0.20498 | 0.12525 | 0.13766 |
| {'Proximal Bilayer_Roughness / Å'} | 3.7412 | 0.50152 | 0.72594 |
| {'PqiABC Coverage'} | 0.056527 | 0.023658 | 0.022898 |
| {'PqiABC roughness'} | 10.381 | 5.3241 | 3.041 |
| {'Distal Bilayer Lipid APM / Å<sup>2</sup>'} | 72.366 | 9.2156 | 9.9243 |
| {'Distal Bilayer coverage'} | 0.58911 | 0.082136 | 0.088965 |
| {'Distal Bilayer position'} | 0.7987 | 0.0076033 | 0.0065961 |
| {'Distal Bilayer HG Additional hyd'} | 0.097508 | 0.066036 | 0.11131 |
| {'Distal Bilayer Roughness / Å'} | 8.9119 | 4.9842 | 4.2571 |
| **Derived Structural Parameters** |  |  |  |
| {'Prox Bilayer Coverage / %'} | 97.377 | 2.0892 | 1.605 |
| {'Prox Bilayer HG thickness / Å'} | 5.5547 | 0.1825 | 0.20559 |
| {'Prox Bilayer Inner Tails thickness / Å'} | 15.261 | 0.5014 | 0.56484 |
| {'Prox Bilayer Outer Tails thickness / Å'} | 15.261 | 0.5014 | 0.56484 |
| {'PqiABC Total Thickness / Å'} | 320 | 0 | 0 |
| {'Bilayer_to_Bilayer Distance / Å'} | 257.07 | 2.3179 | 2.1217 |
| {'Distal Bilayer tails thickness / Å'} | 12.183 | 1.4693 | 1.7777 |
| {'Distal Bilayer HG thickness / Å'} | 4.4343 | 0.5348 | 0.64706 |
| {'Distal Bilayer HG / %'} | 0.05655 | 0.038651 | 0.067257 |

**Supplementary Table 8. Fitted and fitting derived structural parameters from tails deuterated (d) DMPC bilayer with a population of PqiC on its outer surface (data and fits shown in main article Figure 2).**

| Parameter | Mean | Relative_Low_Error | Relative_High_Error |
| --- | --- | --- | --- |
| **Fitted Parameters** |  |  |  |
| {'Substrate_Roughness / Å'} | 7.5334 | 2.8021 | 2.7315 |
| {'SiO2_thickness / Å'} | 8.5257 | 1.9086 | 1.8296 |
| {'SiO2_roughness / Å'} | 3.5011 | 0.97905 | 1.2863 |
| {'SiO2_hydration / %'} | 10.403 | 6.7485 | 8.7823 |
| {'Lipid_APM / Å<sup>2</sup>'} | 50.402 | 2.1497 | 2.1929 |
| {'Proximal Bilayer_Coverage'} | 0.98443 | 0.01797 | 0.010806 |
| {'Proximal Bilayer_HG_Water' } | 0.1373 | 0.087792 | 0.11293 |
| {'Proximal Bilayer_roughness / Å' } | 3.8365 | 0.56926 | 0.78388 |
| {'dDMPC_Tails_Content'} | 0.92354 | 0.019446 | 0.019009 |
| {'Water Thickness / Å'} | 5.0759 | 1.3679 | 1.2339 |
| {'PqiC Coverage'} | 0.12924 | 0.023715 | 0.027019 |
| {'PqiC roughness / Å'} | 5.2866 | 3.3298 | 2.6271 |
| **Fitting Derived Parameters** |  |  |  |
| {'Proximal Bilayer_Coverage / %'} | 98.443 | 1.797 | 1.0806 |
| {'Headgroup Thickness / Å'} | 6.3667 | 0.26551 | 0.28362 |
| {'Tails Thickness / Å'} | 28.852 | 1.2032 | 1.2853 |
| {'Total Thickness / Å'} | 41.585 | 1.7342 | 1.8525 |
| {'Bilayer_dDMPC / %'} | 90.452 | 1.9636 | 2.2241 |
| {'Bilayer_contaminant / %'} | 7.493 | 1.8897 | 1.945 |
| {'PqiC Total Thickness'} | 50 | 0 | 0 |

**Supplementary Table 9. Fitted and fitting derived structural parameters from tails deuterated (d) DMPC bilayer with PqiABC on its outer surface (data and fits shown in main article Figure 2). The formation of the full Pqi complex caused the formation of a small lipid population (distal lipid) around the hydrophobic region of PqiAB.**

| Parameter | Mean | Relative_Low_Error | Relative_High_Error |
| --- | --- | --- | --- |
| **Fitted Parameters** |  |  |  |
| {'Substrate_Roughness / Å' } | 5.8864 | 2.39 | 2.8567 |
| {'SiO2_thickness / Å' } | 9.2087 | 2.2989 | 2.1176 |
| {'SiO2_roughness / Å' } | 3.6521 | 1.0919 | 1.3601 |
| {'SiO2_hydration / %' } | 12.663 | 8.149 | 9.4668 |
| {'Lipid_APM / Å<sup>2</sup>' } | 45.013 | 2.2497 | 2.6461 |
| {'Bilayer_Coverage' } | 0.91464 | 0.03907 | 0.041786 |
| {'Bilayer_HG_Hydration' } | 0.34508 | 0.17229 | 0.1697 |
| {'Bilayer_Roughness_DMPC1 / Å' } | 5.2503 | 1.2569 | 1.2495 |
| {'dDMPC Tails Content Inner leaflet' } | 0.89579 | 0.058841 | 0.053659 |
| {'dDMPC Tails Content Outer leaflet' } | 0.77045 | 0.099347 | 0.08577 |
| {'Water_thickness / Å' } | 3.6217 | 1.6421 | 1.6193 |
| {'PqiABC Coverage' } | 0.17808 | 0.019498 | 0.020019 |
| {'Distal Thickness / Å' } | 18.004 | 8.3376 | 8.1075 |
| {'Distal Total Coverage' } | 0.11054 | 0.04269 | 0.070393 |
| {'Distal Position' } | 0.8018 | 0 | 0 |
| {'PqiABC roughness' } | 9.6572 | 4.1455 | 1.6777 |
| {'Distal d to h lipid mix' } | 0.4039 | 0.19571 | 0.18242 |
| **Fitting Derived Parameters** |  |  |  |
| {'Bilayer Coverage / %' } | 91.464 | 3.907 | 4.1786 |
| {'Bilayer HG Thickness / Å' } | 7.1289 | 0.39576 | 0.37505 |
| {'Bilayer Tails Thickness / Å' } | 32.306 | 1.7934 | 1.6996 |
| {'Bilayer Total Thickness / Å' } | 46.563 | 2.5849 | 2.4497 |
| {'Bilayer Inner Leaflet dDMPC / %' } | 81.792 | 6.3323 | 5.8787 |
| {'Bilayer Inner Leaflet DDM/Native Lipid / %' } | 9.507 | 4.8841 | 5.4047 |
| {'Bilayer Outer Leaflet dDMPC / %' } | 70.302 | 8.4007 | 7.0627 |
| {'Bilayer Outer Leaflet DDM/Native Lipid / %' } | 21.052 | 8.1642 | 9.4561 |
| {'Pqi Total / Å' } | 320 | 0 | 0 |
| {'Bilayer to Distal / Å' } | 270 | 0 | 0 |
| {'Distal Coverage / %' } | 11.054 | 4.269 | 7.0393 |
| {'Distal dDPPC / %' } | 4.2996 | 2.54 | 3.7781 |
| {'Distal Native Lipid / %' } | 6.4007 | 2.8155 | 4.3873 |

**Supplementary Table 10. Fitted and fitting derived structural parameters from double bilayer sample on a PqiABC scaffold with a proximal bilayer deposited as dDMPC and a distal bilayer deposited as hPOPC (data and fits shown in main article Figure 2).**

| Parameter | Mean | Relative_Low_Error | Relative_High_Error |
| --- | --- | --- | --- |
| **Fitted Parameters** |  |  |  |
| {'Substrate_Roughness / Å' } | 5.7498 | 2.4046 | 2.6168 |
| {'SiO2_thickness / Å' } | 9.914 | 2.0664 | 1.7766 |
| {'SiO2_roughness / Å' } | 3.7053 | 1.1024 | 1.4389 |
| {'SiO2_hydration / %' } | 12.377 | 7.4157 | 8.3953 |
| {'Water_thickness / Å' } | 4.1868 | 1.4321 | 1.0529 |
| {'Lipid_APM / Å<sup>2</sup>' } | 52.431 | 3.5469 | 3.2734 |
| {'Bilayer_Coverage' } | 0.97609 | 0.022071 | 0.015996 |
| {'Bilayer_HG_Hydration' } | 0.20861 | 0.13435 | 0.20426 |
| {'Bilayer_Roughness/ Å' } | 3.8279 | 0.56199 | 0.83268 |
| {'Inner_Leaflet_DtoH_Bilayer_Mix_After_POPC' } | 0.65628 | 0.04143 | 0.039775 |
| {'Outer_Leaflet_DtoH_Bilayer_Mix_After_POPC' } | 0.57767 | 0.033554 | 0.031907 |
| {'PqiABC_Coverage' } | 0.15965 | 0.018831 | 0.020406 |
| {'PqiABC_roughness' } | 12.038 | 2.8401 | 1.7972 |
| {'Bilayer 2 Lipid APM / Å<sup>2</sup>' } | 69.751 | 14.146 | 14.417 |
| {'Bilayer 2 coverage' } | 0.58789 | 0.1158 | 0.12656 |
| {'Bilayer 2 DMPC_hLipid_Ratio' } | 0.52591 | 0.027781 | 0.033935 |
| {'Bilayer 2 position' } | 0.81758 | 0.011384 | 0.013331 |
| {'Bilayer 2 HG Additional hyd' } | 0.12354 | 0.079349 | 0.10499 |
| **Fitting Derived Parameters** |  |  |  |
| {'Prox Bilayer Inner leaflet dDMPC / %' } | 65.628 | 4.143 | 3.9775 |
| {'Prox Bilayer Inner leaflet hLipid / %' } | 34.371 | 3.9778 | 4.1423 |
| {'Prox Bilayer Outer leaflet dDMPC / %' } | 57.767 | 3.3554 | 3.1907 |
| {'Prox Bilayer Outer leaflet hLipid / %' } | 42.232 | 3.1906 | 3.3551 |
| {'Prox Bilayer dDMPC Average / %' } | 61.636 | 1.4689 | 1.4967 |
| {'Prox Bilayer hLipid Average / %' } | 38.364 | 1.4967 | 1.4685 |
| {'Prox Bilayer Coverage / %' } | 97.609 | 2.2071 | 1.5996 |
| {'Prox Bilayer HG thickness / Å' } | 6.1203 | 0.35962 | 0.44399 |
| {'Prox Bilayer Inner Tails thickness / Å' } | 14.883 | 0.93134 | 1.1389 |
| {'Prox Bilayer Outer Tails thickness / Å' } | 15.102 | 0.88639 | 1.1182 |
| {'PqiABC Total Thickness / Å' } | 320 | 0 | 0 |
| {'Bilayer_to_Bilayer Distance / Å' } | 263.56 | 3.9535 | 4.1004 |
| {'Dist Bilayer tails thickness / Å' } | 22.976 | 3.8958 | 5.8863 |
| {'Dist Bilayer HG thickness / Å' } | 4.6005 | 0.78815 | 1.1702 |
| {'Dist Bilayer dDMPC Average / %' } | 52.591 | 2.7781 | 3.3935 |
| {'Dist Bilayer hLipid Average / %' } | 47.407 | 3.3935 | 2.7789 |
| {'Dist Bilayer HG / %' } | 0.0721 | 0.047505 | 0.063317 |

**Supplementary Table 11. Fitted and fitting derived structural parameters from tails deuterated (d) DPPC bilayer with a population of PqiC on its outer surface (data and fits shown in main article Figure 3).**

| Parameter | Mean | Relative_Low_Error | Relative_High_Error |
| --- | --- | --- | --- |
| **Fitted Parameters** |  |  |  |
| {'Substrate_Roughness / Å'} | 5.9804 | 2.5002 | 2.6805 |
| {'SiO2_thickness / Å'} | 7.5084 | 1.6094 | 2.0456 |
| {'SiO2_roughness / Å'} | 3.3498 | 0.89464 | 1.3313 |
| {'SiO2_hydration / %'} | 10.865 | 7.1358 | 9.1406 |
| {'Prox Lipid_APM / Å<sup>2</sup>'} | 48.756 | 2.7937 | 4.4743 |
| {'Prox Bilayer_Coverage'} | 0.81627 | 0.04044 | 0.066183 |
| {'Prox Bilayer_HG_Water' } | 0.19777 | 0.13397 | 0.17735 |
| {'Prox Bilayer_roughness / Å'} | 5.3339 | 1.5297 | 1.795 |
| {'Prox dDPPC_Tails_Content'} | 0.8457 | 0.049747 | 0.041237 |
| {'Water Thickness / Å'} | 4.0044 | 1.7893 | 1.8128 |
| {'PqiC Coverage'} | 0.14447 | 0.028227 | 0.029548 |
| {'PqiC roughness / Å'} | 5.0224 | 3.2082 | 2.717 |
| **Fitting Derived Parameters** |  |  |  |
| {'Bilayer_Coverage / %'} | 81.627 | 4.044 | 6.6183 |
| {'Headgroup Thickness / Å'} | 6.5816 | 0.55331 | 0.40001 |
| {'Tails Thickness / Å'} | 33.841 | 2.845 | 2.0568 |
| {'Total Thickness / Å'} | 47.004 | 3.9516 | 2.8568 |
| {'Bilayer_dDPPC / %'} | 68.724 | 4.2895 | 6.0698 |
| {'Bilayer_contaminant (DDM) / %'} | 12.653 | 3.5071 | 4.4552 |
| {'PqiC Total Thickness'} | 50 | 0 | 0 |

**Supplementary Table 12. Fitted and fitting derived structural parameters from tails deuterated (d) DPPC bilayer with PqiABC on its outer surface (data and fits shown in main article Figure 3). The formation of the full Pqi complex caused the formation of a small lipid population (distal lipid) around the hydrophobic region of PqiAB.**

| Parameter | Mean | Relative_Low_Error | Relative_High_Error |
| --- | --- | --- | --- |
| **Fitted Parameters** |  |  |  |
| {'Substrate_Roughness / Å'} | 5.8113 | 2.4573 | 3.1024 |
| {'SiO2_thickness / Å'} | 8.6458 | 2.2532 | 2.6241 |
| {'SiO2_roughness / Å'} | 3.4705 | 0.95019 | 1.4237 |
| {'SiO2_hydration / %'} | 13.395 | 8.1988 | 10.589 |
| {'Lipid_APM / Å<sup>2</sup>'} | 50.74 | 3.1794 | 4.3509 |
| {'Bilayer_Coverage'} | 0.8096 | 0.041569 | 0.062517 |
| {'Bilayer_HG_Hydration'} | 0.2826 | 0.16446 | 0.18322 |
| {'Bilayer_Roughness / Å'} | 5.8872 | 1.8531 | 1.7892 |
| {'dDPPC Tails Content Inner leaflet'} | 0.89458 | 0.067235 | 0.057062 |
| {'dDPPC Tails Content Outer leaflet'} | 0.78126 | 0.09711 | 0.084846 |
| {'Water_thickness / Å'} | 4.3182 | 1.8012 | 1.6071 |
| {'PqiABC Coverage'} | 0.16797 | 0.020315 | 0.022485 |
| {'Distal Thickness / Å'} | 35.049 | 14.854 | 10.449 |
| {'Distal Total Coverage'} | 0.12487 | 0.031778 | 0.06496 |
| {'Distal Position'} | 0.80435 | 0.02408 | 0.025994 |
| {'PqiABC roughness'} | 8.9857 | 4.3968 | 2.1863 |
| {'Distal d to h lipid mix'} | 0.40388 | 0.10432 | 0.090194 |
| **Fitting Derived Parameters** |  |  |  |
| {'Bilayer Coverage / %'} | 80.96 | 4.1569 | 6.2517 |
| {'Bilayer HG Thickness / Å'} | 6.3243 | 0.49947 | 0.42276 |
| {'Bilayer Tails Thickness / Å'} | 35.965 | 2.8404 | 2.4041 |
| {'Bilayer Total Thickness / Å'} | 48.614 | 3.8393 | 3.2497 |
| {'Bilayer Inner Leaflet dDPPC / %'} | 72.494 | 6.3505 | 7.0472 |
| {'Bilayer Inner Leaflet DDM/Native Lipid / %'} | 8.6093 | 4.6703 | 5.5109 |
| {'Bilayer Outer Leaflet dDPPC / %'} | 63.609 | 7.5373 | 6.7135 |
| {'Bilayer Outer Leaflet DDM/Native Lipid / %'} | 17.768 | 7.117 | 8.5626 |
| {'Pqi Total / Å'} | 320 | 0 | 0 |
| {'Bilayer to Distal / Å'} | 258.87 | 7.8243 | 8.4837 |
| {'Distal Coverage / %'} | 12.487 | 3.1778 | 6.496 |
| {'Distal dDPPC / %'} | 5.1476 | 1.9407 | 2.6865 |
| {'Distal Native Lipid / %'} | 7.3981 | 1.9189 | 4.0523 |

**Supplementary Table 13. Fitted and fitting derived structural parameters from double bilayer sample on a PqiABC scaffold with a proximal bilayer deposited as dPPC and a distal bilayer deposited as hPOPC at 20°C (data and fits shown in main article Figures 3 and 4).**

| Parameter | Mean | Relative_Low_Error | Relative_High_Error |
| --- | --- | --- | --- |
| **Fitting Parameters** |  |  |  |
| {'Substrate_Roughness / Å'} | 5.0184 | 1.9002 | 2.4967 |
| {'SiO2_thickness / Å'} | 8.6083 | 2.0567 | 2.3375 |
| {'SiO2_roughness / Å'} | 3.6936 | 1.1495 | 1.4891 |
| {'SiO2_hydration / %'} | 11.732 | 7.7525 | 9.7737 |
| {'Water_thickness / Å'} | 2.5998 | 1.4245 | 1.5685 |
| {'Prox Lipid APM / Å<sup>2</sup>'} | 45.284 | 2.407 | 2.8447 |
| {'Prox Bilayer Coverage'} | 0.95516 | 0.029854 | 0.025566 |
| {'Prox Bilayer HG Hydration'} | 0.37317 | 0.19278 | 0.16006 |
| {'Bilayer_Roughness / Å'} | 5.0523 | 1.3214 | 1.3842 |
| {'Prox Inner_Leaflet_DtoH_Bilayer_Mix_After_POPC' } | 0.68644 | 0.031245 | 0.030803 |
| {'Prox Outer_Leaflet_DtoH_Bilayer_Mix_After_POPC' } | 0.59346 | 0.025879 | 0.026727 |
| {'PqiABC Coverage'} | 0.1757 | 0.021859 | 0.023108 |
| {'PqiABC roughness'} | 12.695 | 1.751 | 0.93316 |
| {'Dist Bilayer Lipid APM / Å<sup>2</sup>'} | 45.658 | 6.6288 | 9.0546 |
| {'Dist Bilayer coverage'} | 0.39855 | 0.036862 | 0.051731 |
| {'Dist Bilayer dPPC_hLipid Ratio'} | 0.20799 | 0.021209 | 0.020507 |
| {'Dist Bilayer position'} | 0.78075 | 0.017517 | 0.019353 |
| {'Dist Bilayer HG Additional hyd'} | 0.15447 | 0.097063 | 0.094741 |
| **Fitting Derived Parameters** |  |  |  |
| {'Prox Bilayer Inner leaflet dPPC / %'} | 68.644 | 3.1245 | 3.0803 |
| {'Prox Bilayer Inner leaflet hLipid / %'} | 31.355 | 3.0805 | 3.1249 |
| {'Prox Bilayer Outer leaflet dDMPC / %'} | 59.346 | 2.5879 | 2.6727 |
| {'Prox Bilayer Outer leaflet hLipid / %'} | 40.653 | 2.6724 | 2.5874 |
| {'Prox Bilayer dPPC Average / %'} | 63.943 | 1.5908 | 1.8591 |
| {'Prox Bilayer hLipid Average / %'} | 36.057 | 1.8591 | 1.5904 |
| {'Prox Bilayer Coverage / %'} | 95.516 | 2.9854 | 2.5566 |
| {'Prox Bilayer HG thickness / Å'} | 7.0861 | 0.41888 | 0.39784 |
| {'Prox Bilayer Inner Tails thickness / Å'} | 18.612 | 1.13 | 1.0718 |
| {'Prox Bilayer Outer Tails thickness / Å'} | 18.723 | 1.1088 | 1.058 |
| {'PqiABC Total Thickness / Å'} | 320 | 0 | 0 |
| {'Bilayer_to_bilayer Distance / Å'} | 251.4 | 5.8752 | 6.0498 |
| {'Dist Bilayer tails thickness / Å'} | 38.093 | 6.2912 | 6.462 |
| {'Dist Bilayer HG thickness / Å'} | 7.0279 | 1.1631 | 1.1937 |
| {'Dist Bilayer dPPC Average / %'} | 20.799 | 2.1209 | 2.0507 |
| {'Dist Bilayer hLipid Average / %'} | 79.2 | 2.0507 | 2.1208 |
| {'Prox Bilayer thickness / Å'} | 51.508 | 3.076 | 2.9267 |
| {'Dist Bilayer thickness / Å'} | 52.276 | 6.2957 | 6.5414 |

**Supplementary Table 14. Fitted and fitting derived structural parameters from double bilayer sample on a PqiABC scaffold with a proximal bilayer deposited as dPPC and a distal bilayer deposited as hPOPC at 42°C (data and fits shown in main article Figure 4).**

| Parameter | Mean | Relative_Low_Error | Relative_High_Error |
| --- | --- | --- | --- |
| **Fitted Parameters** |  |  |  |
| {'Substrate_Roughness / Å'} | 4.877 | 1.9112 | 2.54 |
| {'SiO2_thickness / Å'} | 8.4737 | 1.904 | 2.2836 |
| {'SiO2_roughness / Å'} | 3.5337 | 1.0115 | 1.5793 |
| {'SiO2_hydration / %'} | 14.862 | 9.1002 | 10.13 |
| {'Water_thickness / Å'} | 4.2598 | 1.6419 | 1.1205 |
| {'Lipid_APM / Å<sup>2</sup>'} | 57.104 | 4.3714 | 4.402 |
| {'Prox Bilayer_Coverage'} | 0.97316 | 0.02596 | 0.018492 |
| {'Prox Bilayer_HG_Hydration'} | 0.2479 | 0.16917 | 0.22794 |
| {'Prox Bilayer_Roughness / Å'} | 3.9983 | 0.67863 | 0.91325 |
| {'Prox Inner_Leaflet_DtoH_Bilayer_Mix_After_POPC' } | 0.58179 | 0.033644 | 0.035172 |
| {'Prox Outer_Leaflet_DtoH_Bilayer_Mix_After_POPC' } | 0.5221 | 0.032648 | 0.030903 |
| {'PqiABC Coverage' } | 0.16689 | 0.0213 | 0.019915 |
| {'PqiABC roughness'} | 13.099 | 2.1661 | 1.2781 |
| {'Dist Bilayer Lipid APM / Å<sup>2</sup>'} | 62.644 | 10.237 | 15.842 |
| {'Dist Bilayer coverage'} | 0.64562 | 0.098724 | 0.12388 |
| {'Dist Bilayer DPPC_hLipid Ratio'} | 0.47573 | 0.021142 | 0.02498 |
| {'Dist Bilayer position'} | 0.81172 | 0.014443 | 0.010508 |
| {'Dist Bilayer HG Additional hyd'} | 0.1249 | 0.084645 | 0.11172 |
| **Fitting Derived Parameters** |  |  |  |
| {'Prox Bilayer Inner leaflet dPPC / %'} | 58.179 | 3.3644 | 3.5172 |
| {'Prox Bilayer Inner leaflet hLipid / %'} | 41.818 | 3.516 | 3.3659 |
| {'Prox Bilayer Outer leaflet dPPC / %'} | 52.21 | 3.2648 | 3.0903 |
| {'Prox Bilayer Outer leaflet hLipid / %'} | 47.789 | 3.0903 | 3.2647 |
| {'Prox Bilayer dPPC Average / %'} | 55.231 | 1.4184 | 1.3616 |
| {'Prox Bilayer hLipid Average / %'} | 44.769 | 1.3618 | 1.4185 |
| {'Prox Bilayer Coverage / %'} | 97.316 | 2.596 | 1.8492 |
| {'Prox Bilayer HG thickness / Å'} | 5.6194 | 0.40215 | 0.46586 |
| {'Prox Bilayer Inner Tails thickness / Å' } | 14.864 | 1.0867 | 1.2396 |
| {'Prox Bilayer Outer Tails thickness / Å' } | 14.926 | 1.0738 | 1.2323 |
| {'PqiABC Total Thickness / Å'} | 320 | 0 | 0 |
| {'Bilayer_to_bilayer Distance / Å'} | 261.11 | 4.7106 | 3.5796 |
| {'Dist Bilayer tails thickness / Å'} | 27.29 | 5.5513 | 5.3579 |
| {'Dist Bilayer HG thickness / Å'} | 5.1222 | 1.0339 | 1.0009 |
| {'Dist Bilayer dPPC Average / %'} | 47.573 | 2.1142 | 2.498 |
| {'Dist Bilayer hLipid Average / %'} | 52.427 | 2.498 | 2.1142 |
| {'Prox Bilayer thickness / Å'} | 41.017 | 2.9508 | 3.4287 |
| {'Dist Bilayer thickness / Å'} | 38.436 | 5.2356 | 5.7435 |

**Supplementary Table 15. Fitted and fitting derived structural parameters from tails deuterated (d) DPPC bilayer with a population of PqiC on its outer surface (data and fits shown in supporting information section 2 Figure S4).**

| Parameter | Mean | Relative_Low_Error | Relative_High_Error |
| --- | --- | --- | --- |
| **Fitted Parameters** |  |  |  |
| {'Substrate_Roughness / Å'} | 5.2606 | 2.1345 | 2.5004 |
| {'SiO2_thickness / Å' } | 7.8816 | 1.7479 | 1.7673 |
| {'SiO2_roughness / Å' } | 3.0417 | 0.69826 | 1.1352 |
| {'SiO2_hydration / %' } | 9.4267 | 6.3874 | 8.4616 |
| {'Lipid_APM / Å<sup>2</sup>' } | 46.977 | 2.7199 | 4.8802 |
| {'Bilayer_Coverage' } | 0.75671 | 0.043183 | 0.068895 |
| {'Bilayer_HG_extra_Water' } | 0.23831 | 0.15505 | 0.20801 |
| {'Bilayer_roughness / Å' } | 5.3159 | 1.5202 | 2.4188 |
| {'dDPPC_Tails_Content' } | 0.84614 | 0.05436 | 0.044048 |
| {'Water Thickness / Å' } | 4.3797 | 2.1252 | 2.2834 |
| {'PqiC Coverage' } | 0.16665 | 0.028505 | 0.031947 |
| {'PqiC roughness / Å' } | 5.3051 | 3.237 | 2.5159 |
| **Fitting Derived Parameters** |  |  |  |
| {'Bilayer_Coverage / %' } | 75.671 | 4.3183 | 6.8895 |
| {'Headgroup Thickness / Å'} | 6.8309 | 0.64286 | 0.41975 |
| {'Tails Thickness / Å' } | 35.123 | 3.3055 | 2.1583 |
| {'Total Thickness / Å' } | 48.785 | 4.5912 | 2.9978 |
| {'Bilayer_dDPPC / %' } | 63.313 | 3.9063 | 6.8153 |
| {'Bilayer_contaminant / %'} | 11.724 | 3.5108 | 4.6075 |
| {'PqiC Total Thickness' } | 50 | 0 | 0 |

**Supplementary Table 16. Fitted and fitting derived structural parameters from tails deuterated (d) DPPC bilayer with PqiABC on its outer surface (data and fits shown in supporting information section 2 Figure S4).**

| Parameter | Mean | Relative_Low_Error | Relative_High_Error |
| --- | --- | --- | --- |
| **Fitted Parameters** |  |  |  |
| {'Substrate_Roughness / Å' } | 5.0363 | 2.0185 | 2.7211 |
| {'SiO2_thickness / Å' } | 9.4107 | 2.1379 | 1.7955 |
| {'SiO2_roughness / Å' } | 3.1003 | 0.7471 | 1.0986 |
| {'SiO2_hydration / %' } | 10.374 | 6.6668 | 9.1384 |
| {'Prox Lipid_APM / Å<sup>2</sup>' } | 44.956 | 2.2158 | 2.4843 |
| {'Prox Bilayer_Coverage' } | 0.73932 | 0.027955 | 0.041998 |
| {'Prox Bilayer_HG_Hydration' } | 0.12481 | 0.086011 | 0.12782 |
| {'Prox Bilayer_Roughness / Å' } | 5.5714 | 1.5329 | 1.5928 |
| {'Prox dDPPC Tails Content Inner leaflet' } | 0.85503 | 0.067566 | 0.05467 |
| {'Prox dDPPC Tails Content Outer leaflet' } | 0.71068 | 0.086673 | 0.074928 |
| {'Water_thickness / Å' } | 5.1643 | 1.5286 | 1.1301 |
| {'PqiABC Coverage' } | 0.15129 | 0.027722 | 0.031482 |
| {'Amphipol Thickness / Å' } | 32.532 | 8.5434 | 10.674 |
| {'Amphipol Coverage' } | 0.006633 | 0.0047408 | 0.010302 |
| {'Distal Position' } | 0.8444 | 0 | 0 |
| {'PqiABC roughness' } | 10.614 | 6.3312 | 4.3607 |
| **Fitting Derived Parameters** |  |  |  |
| {'Prox Bilayer Coverage / %' } | 73.932 | 2.7955 | 4.1998 |
| {'Prox Bilayer HG Thickness / Å' } | 7.138 | 0.3738 | 0.36995 |
| {'Prox Bilayer Tails Thickness / Å' } | 40.592 | 2.1257 | 2.1039 |
| {'Prox Bilayer Total Thickness / Å' } | 54.868 | 2.8734 | 2.8438 |
| {'Bilayer Inner Leaflet dDPPC / %' } | 63.485 | 5.7117 | 5.3514 |
| {'Bilayer Inner Leaflet DDM/Native Lipid /Protein %' } | 10.793 | 4.0652 | 5.078 |
| {'Bilayer Outer Leaflet dDPPC / %' } | 52.746 | 6.3512 | 5.7692 |
| {'Bilayer Outer Leaflet DDM/Native Lipid /Protein %' } | 21.55 | 5.8132 | 6.8007 |
| {'Pqi Total / Å' } | 320 | 0 | 0 |
| {'Bilayer to Distal / Å' } | 275 | 0 | 0 |
| {'Amphipol Coverage / %' } | 0.6633 | 0.47408 | 1.0302 |

**Supplementary Table 17. FFitted and fitting derived structural parameters from double bilayer sample on a PqiABC scaffold with a proximal bilayer deposited as dPPC and a distal bilayer deposited as hPOPC at 28°C (data and fits shown in supporting information section 2 Figures S4 and S5).**

| Parameter | Mean | Relative_Low_Error | Relative_High_Error |
| --- | --- | --- | --- |
| **Fitted Parameters** |  |  |  |
| {Substrate_Roughness / Å} | 5.1149 | 2.0739 | 2.5092 |
| {SiO<sub>2</sub>_thickness / Å} | 8.8075 | 2.2227 | 2.0888 |
| {SiO<sub>2</sub>_roughness / Å} | 3.4468 | 0.97959 | 1.265 |
| {SiO<sub>2</sub>_hydration / %} | 13.115 | 7.5746 | 8.7294 |
| {Water_thickness / Å} | 3.137 | 1.5753 | 1.5096 |
| {Prox Lipid_APM / Å<sup>2</sup>} | 46.406 | 2.784 | 2.8745 |
| {Prox Bilayer_Coverage'} | 0.96131 | 0.029049 | 0.024765 |
| {Prox Bilayer_HG_Hydration'} | 0.29385 | 0.17399 | 0.19032 |
| {Prox Bilayer_Roughness / Å} | 5.21 | 1.3123 | 1.3346 |
| {Prox Inner_Leaflet_D/H_Bilayer_Mix_After_PPPC'} | 0.63825 | 0.028853 | 0.030068 |
| {Prox Outer_Leaflet_D/H_Bilayer_Mix_After_PPPC'} | 0.51427 | 0.0239 | 0.028529 |
| {PqiABC Coverage'} | 0.16457 | 0.022667 | 0.024299 |
| {PqiABC roughness'} | 11.172 | 3.5191 | 1.8053 |
| {Dist Lipid APM / Å<sup>2</sup>} | 68.656 | 14.296 | 13.319 |
| {Dist Bilayer coverage'} | 0.53458 | 0.11117 | 0.094587 |
| {Dist Bilayer D/H Ratio'} | 0.26514 | 0.021329 | 0.021475 |
| {Dist Bilayer position'} | 0.81316 | 0.016485 | 0.013636 |
| {Dist Bilayer HG Additional hyd'} | 0.16225 | 0.10045 | 0.090618 |
| **Fitting derived Parameters** |  |  |  |
| {Prox Bilayer Inner leaflet dPPC / %} | 63.825 | 2.8853 | 3.0068 |
| {Prox Bilayer Inner leaflet hLipid / %} | 36.174 | 3.0068 | 2.885 |
| {Prox Bilayer Outer leaflet dPPC / %} | 51.427 | 2.39 | 2.8529 |
| {Prox Bilayer Outer leaflet hLipid / %} | 48.572 | 2.853 | 2.39 |
| {Prox Bilayer dPPC Average / %} | 57.808 | 1.4852 | 1.4177 |
| {Prox Bilayer hLipid Average / %} | 42.192 | 1.4177 | 1.4849 |
| {Prox Bilayer Coverage / %} | 96.131 | 2.9049 | 2.4765 |
| {Prox Bilayer HG thickness / Å} | 6.9149 | 0.40337 | 0.44127 |
| {Prox Bilayer Inner Tails thickness / Å} | 18.225 | 1.1009 | 1.1768 |
| {Prox Bilayer Outer Tails thickness / Å} | 18.366 | 1.0705 | 1.1772 |
| {PqiABC Total Thickness / Å} | 320 | 0 | 0 |
| {Bilayer_to_bilayer Distance / Å} | 261.88 | 5.203 | 4.1879 |
| {Dist Bilayer tails thickness / Å} | 25.248 | 4.1049 | 6.6364 |
| {Dist Bilayer HG thickness / Å} | 4.6738 | 0.75934 | 1.2291 |
| {Dist Bilayer dPPC Average / %} | 26.514 | 2.1329 | 2.1475 |
| {Dist Bilayer hLipid Average / %} | 73.485 | 2.1473 | 2.1331 |
| {Dist Bilayer HG / %} | 0.082937 | 0.052312 | 0.054367 |
| {Prox Bilayer thickness / Å} | 50.424 | 2.9721 | 3.2391 |
| {Dist Bilayer thickness / Å} | 39.067 | 4.0527 | 6.76 |

**Supplementary Table 18. Fitted and fitting derived structural parameters from double bilayer sample on a PqiABC scaffold with a proximal bilayer deposited as dPPC and a distal bilayer deposited as hPOPC at 42°C (data and fits shown in supporting information section 2 Figure S5).**

| Parameter | Mean | Relative_Low_Error | Relative_High_Error |
| --- | --- | --- | --- |
| **Fitted Parameters** |  |  |  |
| {'Substrate_Roughness / Å'} | 4.9914 | 2.016 | 2.3818 |
| {'SiO2_thickness / Å'} | 8.394 | 1.8392 | 1.8269 |
| {'SiO2_roughness / Å'} | 3.3806 | 0.92343 | 1.233 |
| {'SiO2_hydration / %'} | 12.728 | 7.9683 | 9.2037 |
| {'Water_thickness / Å'} | 3.3376 | 1.6355 | 1.4195 |
| {'Prox Lipid_APM / Å<sup>2</sup>'} | 53.788 | 4.0126 | 4.1021 |
| {'Prox Bilayer_Coverage'} | 0.97832 | 0.025605 | 0.015194 |
| {'Prox Bilayer_HG_Hydration'} | 0.25432 | 0.16934 | 0.23554 |
| {'Prox Bilayer_Roughness/ Å'} | 4.1354 | 0.75478 | 1.1678 |
| {'Prox Inner_Leaflet_D/H_Bilayer_Mix_After_POPC'} | 0.55915 | 0.029684 | 0.031871 |
| {'Prox Outer_Leaflet_D/H_Bilayer_Mix_After_POPC'} | 0.48896 | 0.025445 | 0.028132 |
| {'PqiABC Coverage'} | 0.1642 | 0.022172 | 0.021819 |
| {'PqiABC roughness'} | 11.037 | 2.881 | 1.8116 |
| {'Dist Bilayer Lipid APM / Å'} | 72.449 | 12.943 | 10.842 |
| {'Dist Bilayer coverage'} | 0.59518 | 0.10896 | 0.07541 |
| {'Dist Bilayer 2 D/H Ratio'} | 0.47466 | 0.023788 | 0.024479 |
| {'Dist Bilayer position'} | 0.81359 | 0.012079 | 0.0081111 |
| {'Dist Bilayer HG Additional hyd'} | 0.14539 | 0.095143 | 0.09801 |
| **Fitting Derived Parameters** |  |  |  |
| {'Prox Bilayer Inner leaflet dPPC / %'} | 55.915 | 2.9684 | 3.1871 |
| {'Prox Bilayer Inner leaflet hLipid / %'} | 44.084 | 3.1874 | 2.9681 |
| {'Prox Bilayer Outer leaflet dPPC / %'} | 48.896 | 2.5445 | 2.8132 |
| {'Prox Bilayer Outer leaflet hLipid / %'} | 51.103 | 2.8132 | 2.5445 |
| {'Prox Bilayer dPPC Average / %'} | 52.473 | 1.1792 | 1.2698 |
| {'Prox Bilayer hLipid Average / %'} | 47.527 | 1.27 | 1.1792 |
| {'Prox Bilayer Coverage / %'} | 97.832 | 2.5605 | 1.5194 |
| {'Prox Bilayer HG thickness / Å'} | 5.9659 | 0.42273 | 0.48093 |
| {'Prox Bilayer Inner Tails thickness / Å'} | 15.798 | 1.1284 | 1.2987 |
| {'Prox Bilayer Outer Tails thickness / Å'} | 15.878 | 1.1341 | 1.2719 |
| {'PqiABC Total Thickness / Å'} | 320 | 0 | 0 |
| {'Bilayer_to_bilayer Distance / Å'} | 261.91 | 4.0034 | 2.6582 |
| {'Dist Bilayer tails thickness / Å'} | 23.588 | 3.0807 | 5.1585 |
| {'Dist Bilayer HG thickness / Å'} | 4.4292 | 0.57658 | 0.96329 |
| {'Dist Bilayer dPPC Average / %'} | 47.466 | 2.3788 | 2.4479 |
| {'Dist Bilayer hLipid Average / %'} | 52.534 | 2.4482 | 2.3788 |
| {'Dist Bilayer HG / %'} | 0.082695 | 0.054478 | 0.060974 |
| {'Prox Bilayer thickness / Å'} | 43.606 | 3.1026 | 3.5286 |
| {'Dist Bilayer thickness / Å'} | 35.443 | 2.9096 | 5.3939 |
